# Exploring structural diversity across the protein universe with The Encyclopedia of Domains

**DOI:** 10.1101/2024.03.18.585509

**Authors:** A. M. Lau, N. Bordin, S. M. Kandathil, I. Sillitoe, V. P. Waman, J. Wells, C. A. Orengo, D. T. Jones

## Abstract

The AlphaFold Protein Structure Database (AFDB) contains full-length predictions of the three-dimensional structures of almost every protein in UniProt. Because protein function is closely linked to structure, the AFDB is poised to revolutionise our understanding of biology, evolution and more. Protein structures are composed of domains, independently folding units that can be found in multiple structural contexts and functional roles. The AFDB’s potential remains untapped due to the difficulty of characterising 200 million structures. Here we present The Encyclopedia of Domains or TED, which combines state-of-the-art deep learning-based domain parsing and structure comparison algorithms to segment and classify domains across the whole AFDB. TED describes over 370 million domains, over 100 million more than detectable by sequence-based methods. Nearly 80% of TED domains share similarities to known superfamilies in CATH, greatly expanding the set of known protein structural domains. We uncover over 10,000 previously unseen structural interactions between superfamilies, expand domain coverage to over 1 million taxa, and unveil thousands of architectures and folds across the unexplored continuum of protein fold space. We expect TED to be a valuable resource that provides a functional interface to the AFDB, empowering it to be useful for a multitude of downstream analyses.

## Introduction

The AlphaFold Protein Structure Database (AFDB)^1,2^ is a groundbreaking initiative which significantly broadened the protein structure universe by expanding 3D representation to over 200 million UniProt sequences. The implications of the AFDB have been profound, not only in academic research in the life sciences but also in the commercial sphere, where the integration of novel data and the advanced technologies used to generate accurate structures is being explored for next-generation structure-based drug discovery^3^.

Notwithstanding its revolutionary impact, the AFDB is not without its limitations and presents a new set of challenges. The sheer scale of the data makes many traditional tools and pipelines, originally designed for considerably smaller datasets, inadequate for navigating the extensive number of structures and sequences within. This necessitates an evolved strategy, warranting a new perspective on how best to represent and traverse the data and complex relationships within such an expansive database, as well as requiring synergy between new algorithmic methods and computational hardware. Recent studies explored AFDB by partitioning full-length AFDB models into clusters of structurally similar proteins, as well as characterising their functions^4,5^. At a more granular level, Bordin et al.^6^ and Schaeffer et al.^7^ interrogated the composition of specific proteomes (the initial AFDB 21 model organism dataset, and 48 proteomes, respectively), cataloguing domains under the CATH^8,9^ and ECOD^10^ frameworks.

Domain discovery is possible via sequence- and structure-based approaches, with Pfam^11,12^ and Gene3D^13^ as prime examples of the former. The Pfam database describes collections of protein families, each represented by a multiple sequence alignment (MSA) and a hidden Markov model (HMM) profile^11,12^. In Gene3D, sequences of existing CATH superfamilies, assigned via structure, are leveraged to discover new domains in sequence space via representative profile HMMs^13^. Sequence-based discovery allows for greater coverage but is limited by HMMs detection capabilities, often failing on remote relatives. Structure-based assignments allow for higher quality domain boundaries and can reveal very remote relatives but have been limited by the low numbers of experimental structures.

An essential perspective yet to be explored in the context of the AFDB is the comprehensive mapping and analysis of protein domains across various branches of the Tree of Life using structural data. The relationships between protein folds and domains have been extensively highlighted by structural databases such as CATH^8,14,15^, ECOD^10^, SCOP^16^ and SCOPe^17^, with differences in criteria for which regions are assigned to domains in proteins. Although early comparisons across domain databases showed agreement for fold level assignments^10,18,19^, significant differences exist for the definition of a protein domain. CATH recognises that some structures can be decomposed into further structural units, each with their own repertoire of observed variations, while SCOP takes into account the idea of fold recurrence - whether a subunit has been observed as reoccuring in another family or only as a single domain^18^.

Examination of the AFDB through the lens of the CATH framework holds the potential to reveal unprecedented insights that can illuminate subtle yet important connections between structure and function across large swathes of organisms. As such, a structure-based mapping of domains within the AFDB promises not only to massively facilitate the exploration of these relationships, but also provide a foundation for further annotation.

In this study, we present a comprehensive analysis of domain composition for the entirety of the AFDB (version 4). This description encompasses over 371 million putative domains, derived from more than 214 million protein sequences across more than 1 million taxa. The identification of these domain structures is made via the consensus of three automated parsing methodologies: Merizo^20^, Chainsaw^21^ and UniDoc^22^. Further, by employing structural comparison methods such as Foldseek and an in-house deep learning method called Merizo-search, more than 251 million domains can be placed on the CATH hierarchy.

## Results

### A high-throughput strategy for identifying structural domains in the AFDB

The AFDB V4 contains predicted structures for over 214 million UniProt sequences across more than 1 million unique taxa, with protein lengths up to 2700 residues (Supp. Figure 1). The first step of our analysis workflow involves removing structures of identical sequences, producing a set of 188 million non-redundant sequences across 653,460 taxa, which we refer to as TED-100.

Our workflow combines three state-of-the-art domain parsing methods together with structure classification algorithms to identify known domain folds (Figure 1a-b; Methods) within the AFDB. Using this workflow, we identified a total of 371 million ‘TED’ domains across the AFDB (Supp. Figure 2, Supp. Figure 3) - 100 million more domains than found via sequence-based methods (Figure 1c). TED-100 describes a roughly 42:55 division between single and multidomain proteins (Figure 1d-i), with the latter up to 20 domains in composition (Supp. Figure 4). Only 2.8% of targets (5.3 million) in TED-100 lack identifiable domains, compared to 33.9% (64.1 million targets) in Gene3D and 26.2% in Pfam (49.4 million targets) (Supp. Figure 5). In TED, these targets either consist entirely of non-domain residues (NDR) or lack any consensus among the three domain segmentation methods employed. Across superkingdoms, the fraction of NDRs varies, with approximately 10% identified across archaeal and bacterial lineages and up to 30% in eukaryotes (Supp. Figure 6).

**Figure 1.**
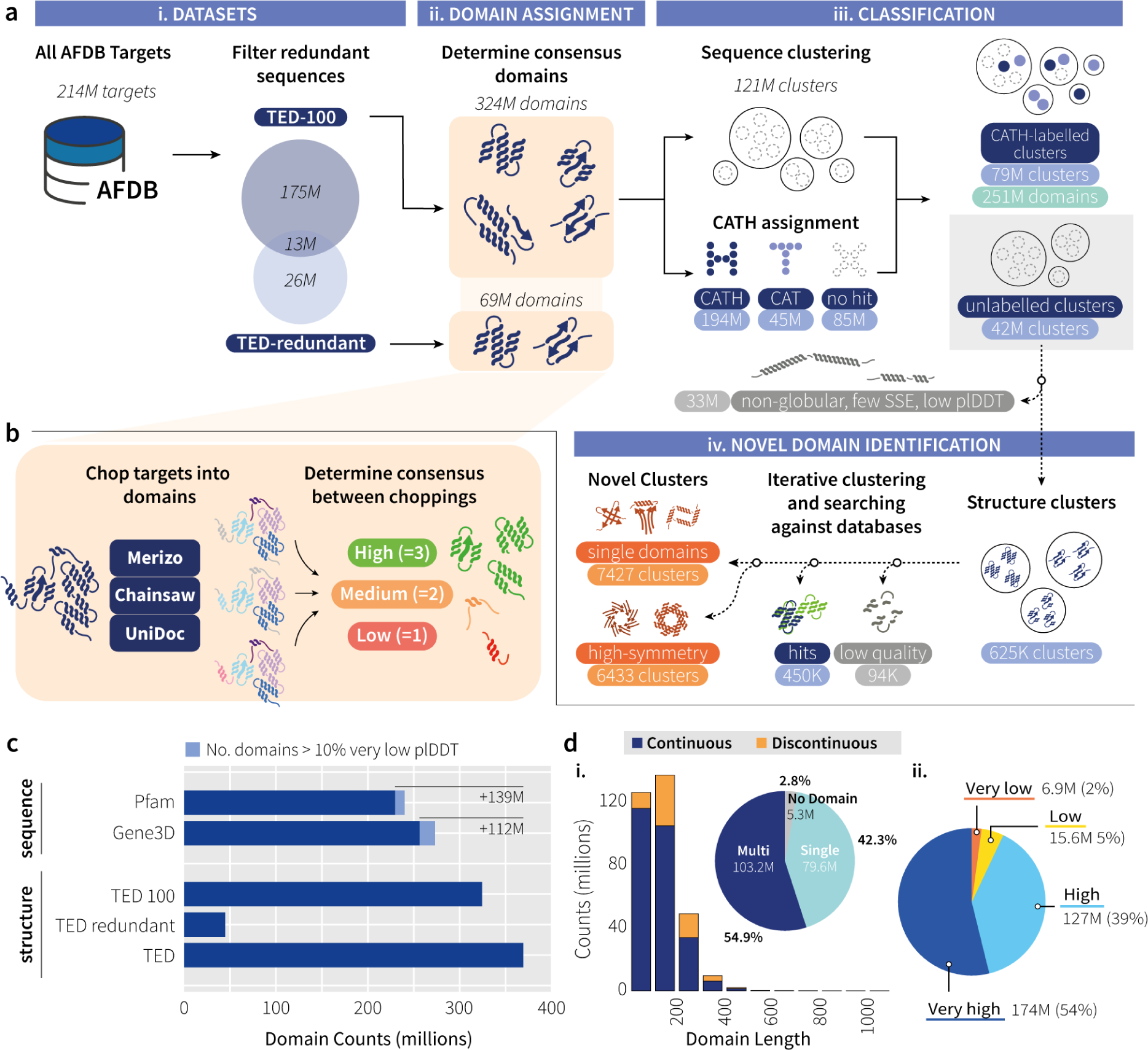
Overall workflow. **a**) i. 214 million AFDB target sequences are filtered by 100% sequence identifying 188 million non-redundant targets (TED-100) and a set of redundant targets (TED-redundant). ii. Both TED-100 and TED-redundant undergo automated domain parsing, with assignments derived from consensus among the three methods. iii. TED-100 domains are processed by MMseqs2, creating over 121 million clusters at 50% identity. Concurrently, domains are matched to CATH domains via Foldseek and Merizo-search, categorised into superfamily (C.A.T.H), topology (C.A.T), or no-matches. Domains found by Merizo-search nearest neighbour matches are considered as topology-level matches. Clusters are annotated with CATH labels, creating partially labelled and unlabelled clusters. Low-quality domains in unlabeled clusters are filtered out. iv. Resultant domains undergo a new workflow for identification, involving clustering and database searches for matches to known structures. Poor quality domains (non-protein-like) are identified using an in-house deep learning method (Methods). Novel domains are additionally scored on internal symmetry using the SymD program (Methods). **b**) Full-length targets are subjected to automated domain parsing by Merizo, Chainsaw and UniDoc. A consensus is taken by identifying assignments where three (high), two (medium) methods agree or no consensus is found (low). Only high and medium consensus domains are analysed further. **c**) Comparison of domains identified by sequence (Pfam and Gene3D) versus structure-based methods (TED). The "TED" count combines TED-100 and TED-redundant, including the 13 million overlapping targets. **d**) i. Domain length distribution and proportion of identified continuous (blue) and discontinuous (orange) domains. Inset shows proportion of single, multi-domain and number of targets with no identified domains (n=188,914,411). ii. Average plDDT distribution for TED-100 domains (n=324,389,697) across confidence bins: dark blue/very high (plDDT >= 90), blue/high (90 > plDDT >= 70), yellow/ low (70 > plDDT >= 50), and orange/very low (plDDT < 50).

Analysis of the average plDDT scores for TED-100 reveals that the majority of domains fall within the "very high" to "high" bins, with only approximately 2% within the lowest bin (Figure 1d-ii). Since our domain segmentation methods do not consider residue plDDT when determining domains, this suggests that our domain finding pipeline effectively identifies plausible domains within the well-folded areas of AF2 models.

### Classification of TED domains into the CATH hierarchy

The 324 million domains in TED-100 were clustered by sequence using MMseqs2^23^ and compared against CATH representative domains using fast structure searching methods (Figure 1a-iii, Methods). This produced approximately 121 million clusters at 50% sequence identity and a minimum coverage of 90% (Supp. Figure 7; Methods). The majority of these clusters comprise just singleton sequences (roughly 81 million), with the largest non-singleton cluster comprising 12,847 domains.

Parallel to sequence clustering, we use a combination of Foldseek^24^ and Merizo-search^25^ (an in-house structure search method using domain coordinate embedding) to search all TED-100 domains against CATH SSG5 domains^9^ (Methods), allowing 194 million domains to be assigned with CATH superfamily (H) labels and 36 million at the topology (T) level (Methods). These labels were further validated by scanning domain sequences against an updated library of HMMs for CATH PDB domains. Approximately 171 million superfamily predictions by Foldseek could be confirmed with exact HMM superfamily matches (88.54%), with an additional 1.8 million domains (0.95%) confirmable at the fold level (0.95%). Only 4.1 million of the 16 million Foldseek predictions for fold matches on CATH can be validated by HMM scans (25.8%), with 11.8 million fold predictions and 20.3 million superfamily predictions by Foldseek not confirmed by an HMM match, suggesting an expansion by 15.3% in CATH labelled domain coverage using AFDB structures over HMM-based sequence assignments.

By identifying sequence clusters with any CATH label assigned domains, the clusters were partitioned into two categories: 78 million clusters (over 251 million domains, nearly 80% of all TED-100 domains) which contain at least one CATH-labelled member, spanning 148 million proteins, and 26 million proteins having no domain annotations in Pfam and and 30 million none in Gene3D (Supp. Figure 8). The remaining 41 million clusters have no members with CATH labels (approximately 73 million domains; Supp. Figure 7). The absence of similarity to CATH domains in the latter clusters could be attributed to them being novel folds, extremely divergent relatives of existing CATH domains or simply being incorrect models, making them unmatchable to known folds.

### Enrichment of Fold Representation by TED

To develop an understanding of how folds are distributed across the AFDB, we assessed the composition of TED using the CATH hierarchy. Figure 2a shows the top 100 CATH superfamilies of each class (alpha, beta and alpha/beta), which are greatly enriched in TED-100 compared to baseline sequence hits in Gene3D.

**Figure 2.**
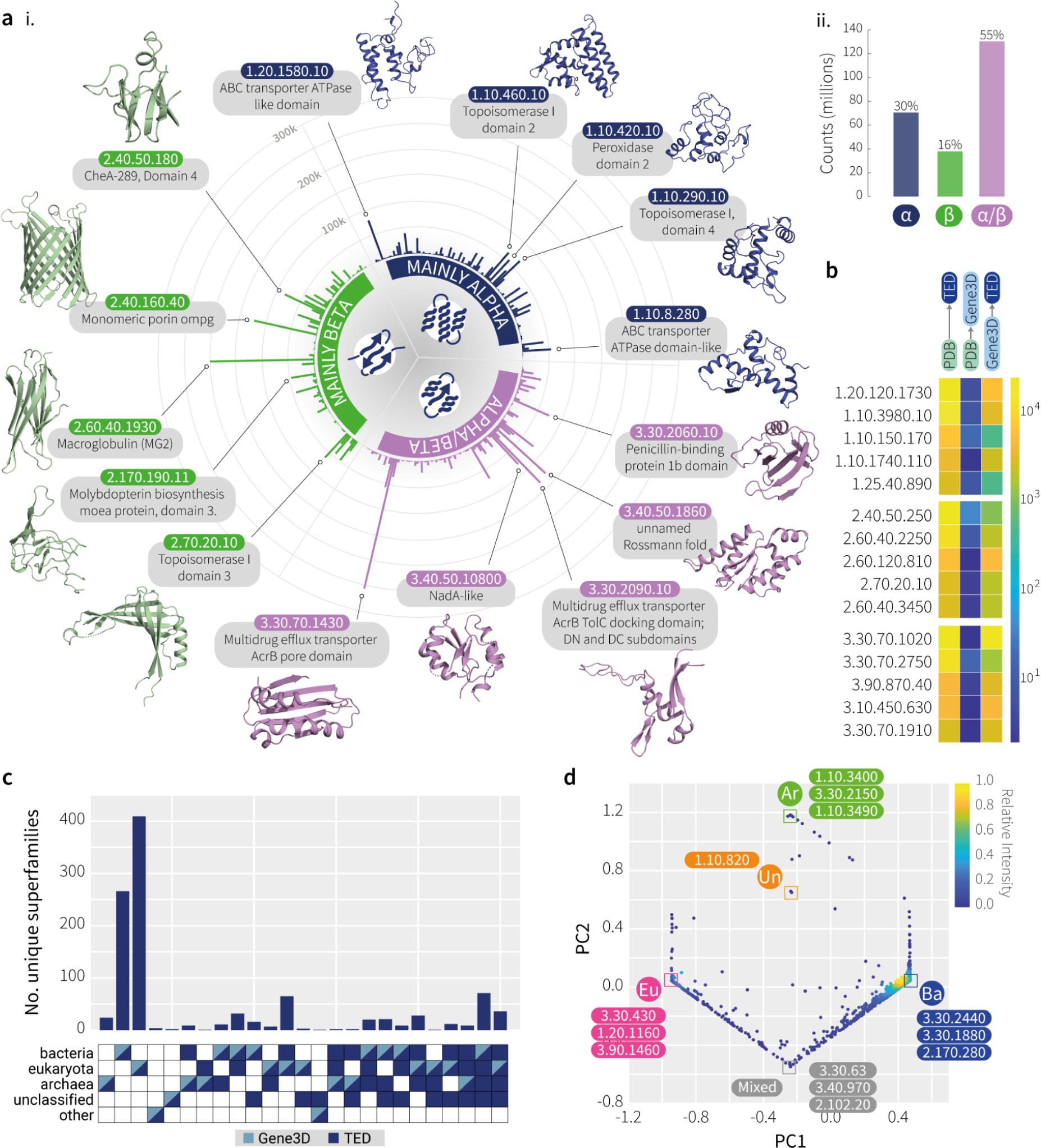
Classification of TED domains using the CATH hierarchy. **(a)** i. The top 100 superfamilies in TED-100 for each CATH class where more matches to CATH superfamilies have been identified via structural hits in TED, compared to sequence hits in Gene3D. ii. Proportion of domains matched to CATH classes (n=238,569,631). **(b)** Enrichment of superfamily representation in TED-100 compared to PDB and Gene3D. The top 5 superfamilies of each CATH class are shown, where enrichment in TED-100 compared to PDB is the greatest. Colour scale represents fold-change in superfamily representation in PDB and Gene3D compared to Gene3D and TED. (**c**) Expansion of CATH superfamilies to new superkingdoms in TED. Plot shows the number of unique superfamilies found in each superkingdom (across the 653,460 taxa of TED-100) according to Gene3D and TED assignments. Only superfamilies where Gene3D domains are exclusive to a single superkingdom are shown (n=1061). **(d)** Exclusivity of CATH topologies across superkingdoms. PCA of normalised CATH topology counts across five superkingdoms: eukaryota (Eu), bacteria (Ba), archaea (Ar), unclassified and other sequences (Un). The ‘mixed’ category comprises topologies found in roughly equal proportions in Eu/Ba domains. Examples of superkingdom-exclusive topologies are shown for each category.

These folds and architectures are unevenly distributed across the Tree of Life, with 0.5%, 9% and 11% of CATH folds being exclusively found in the archaea, eukarya and bacteria. 61% of CATH folds are reused across all superkingdoms, suggesting essential roles for cellular life, with the remaining 18.5% being found in only two superkingdoms with shared evolutionary trajectories.

The most abundant superfamilies in TED-100 and CATH are shown in Supp. Figure 9 and Supp. Figure 10. Directly comparing the top superfamilies assigned in TED compared to CATH, in terms of raw domain counts, sees the promotion of several superfamilies into the top 5 of each class, including the MFS general substrate transporter-like domain, translation factors and the FAD/NAD(P)-binding domain (Supp. Figure 9). The set of superfamilies highly enriched in TED include those associated with the archetypal multi-drug efflux pump AcrB. AcrB forms part of the AcrAB-TolC efflux pump in bacteria where it is responsible for the export of harmful substances such as antibiotics, contributing to antibiotic resistance^26^. The constituent domains of AcrB, including the pore domain, transmembrane domain and TolC docking domain, are found greatly enriched in TED compared to the PDB, providing up to nearly a thousand-fold increase in representation for these biologically important superfamilies.

The pore domain, which forms part of the pore selectivity filter in RND transporters, is principally found in bacterial species and in a small minority of archaeal and eukaryotic lineages based on Gene3D hits. However, structure-based searching via TED expands the coverage of the pore domain superfamily into an additional 18 archaeal, 1315 bacterial and 284 eukaryotic lineages unique to TED, which evaded even HMM searches. This broader coverage of organisms, revealed by structural comparisons, may reveal potential evolutionary events, such as lateral gene transfer between bacteria and eukaryotes. This expansion is exemplified in Figure 2c, where we show that over 1000 CATH superfamilies, described by Gene3D sequence matches as occurring solely in a single superkingdom, could actually be mapped to other branches when structure was taken into account in TED.

TED also enables us to better study the distribution of folds across different branches of the tree of life. Among the 193 million TED domains with superfamily labels, we observed folds that were exclusively localised to specific superkingdoms. These findings are shown in Figure 2d, where folds are summarised at the CATH topology level, visualised by principal component analysis (PCA), and where points at the vertices represent exclusive occurrence of a topology and all its superfamilies, in a particular superkingdom.

### Novel high-symmetry architectures

From our TED workflow (Figure 1), we identified 41 million sequence clusters which could not be linked to CATH superfamilies. Representatives of these clusters were subjected to a workflow aimed at identifying novel domain folds (Figure 1a-iv; Methods).

While reviewing these clusters, it was apparent that we would need to treat repeat architectures with high internal symmetry separately. A good example of domains in this class are the various WD40 beta propellers, which are considered distinct domain architectures in their own right, but clearly comprise repeats of domain-like units. To identify similar domains in our workflow, we calculated Z-scores using the SymD program^27^, sequestering any cluster representatives with a symmetry Z-score of greater than 9, into a new category of 6,433 highly symmetric novel fold clusters (Supp. Figure 11).

Within these clusters, we find new architectures such as an 11-bladed beta-propeller and an 11-helix propeller which have not been seen before (Figure 3). Various other propeller arrangements including the “beta-flower” domain shown by Durairaj et al.^4^, are also identified, with some visually striking examples shown in Figure 3. More curiously, we find a broad new category of architectures which are composed of cyclic repeats, extruded along an axis to form a highly repetitive and symmetric projection, which we call “extruded repeats”. These domains are highly varied and may be embellished with a variety of secondary structure decorations around the extruded core of the domain.

**Figure 3.**
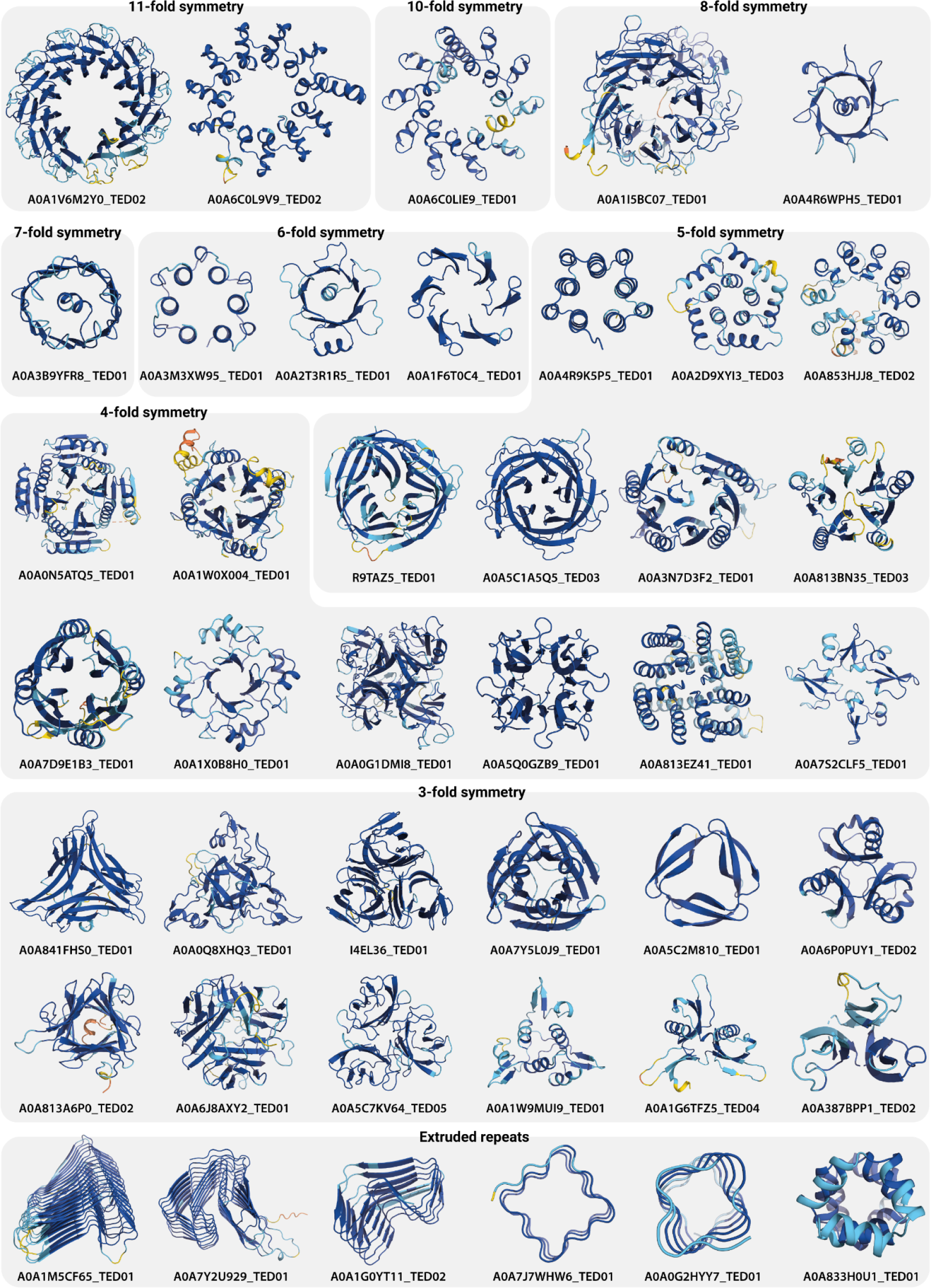
Examples of high-symmetry domains and extruded repeats. Domains are identified as part of the novel domain identification pipeline and are identified as domains with high internal symmetry via scoring with the SymD program (Methods). Extruded repeats are domains with a high number of ordered cyclical repeats projecting along one axis. Colouration follows plDDT confidence bins as per the AFDB (dark blue/very high: plDDT >= 90, blue/high: 90 > plDDT >= 70, yellow/low: 70 > plDDT >= 50 and orange/very low: plDDT < 50).

### Novel Domains and their distribution across the Tree of Life

The remaining low-symmetry clusters were assessed on domain quality, using a variant of the Foldclass network^25^ trained to identify poor quality domain choppings (Supp. Methods), and novelty by further matching against known structure libraries. The final output of our workflow produced 7,427 clusters of putative domains which appear to be well-folded but dissimilar to any known domain fold. Although there is no exact boundary between novel and just highly divergent examples of known folds, by applying a density-based anomaly detection algorithm (Methods), we could at least rank these domains by novelty relative to known domains.

Figure 4 showcases examples of novel domains identified by our workflow. Over a quarter of clusters corresponded to singletons at the sequence cluster level (1930 domains), which are uniformly distributed in terms of the Foldclass novelty score. Most novel domains identified are from bacterial proteins, which is unsurprising given that the latter comprise most proteins in the AFDB and in our TED-100 dataset. Overall, these domains are distributed evenly across different phyla, but several bacterial phyla were found overrepresented when compared to baseline counts across all domains in TED-100, specifically in the PVC group, Myxococcota, Spirochaetota, Bdellovibrionota, Nitrospinae/Tectomicrobia group and Calditrichota (Supp. Figure 12), suggesting that species within these phyla may be underrepresented in terms of domain coverage in the current PDB.

**Figure 4.**
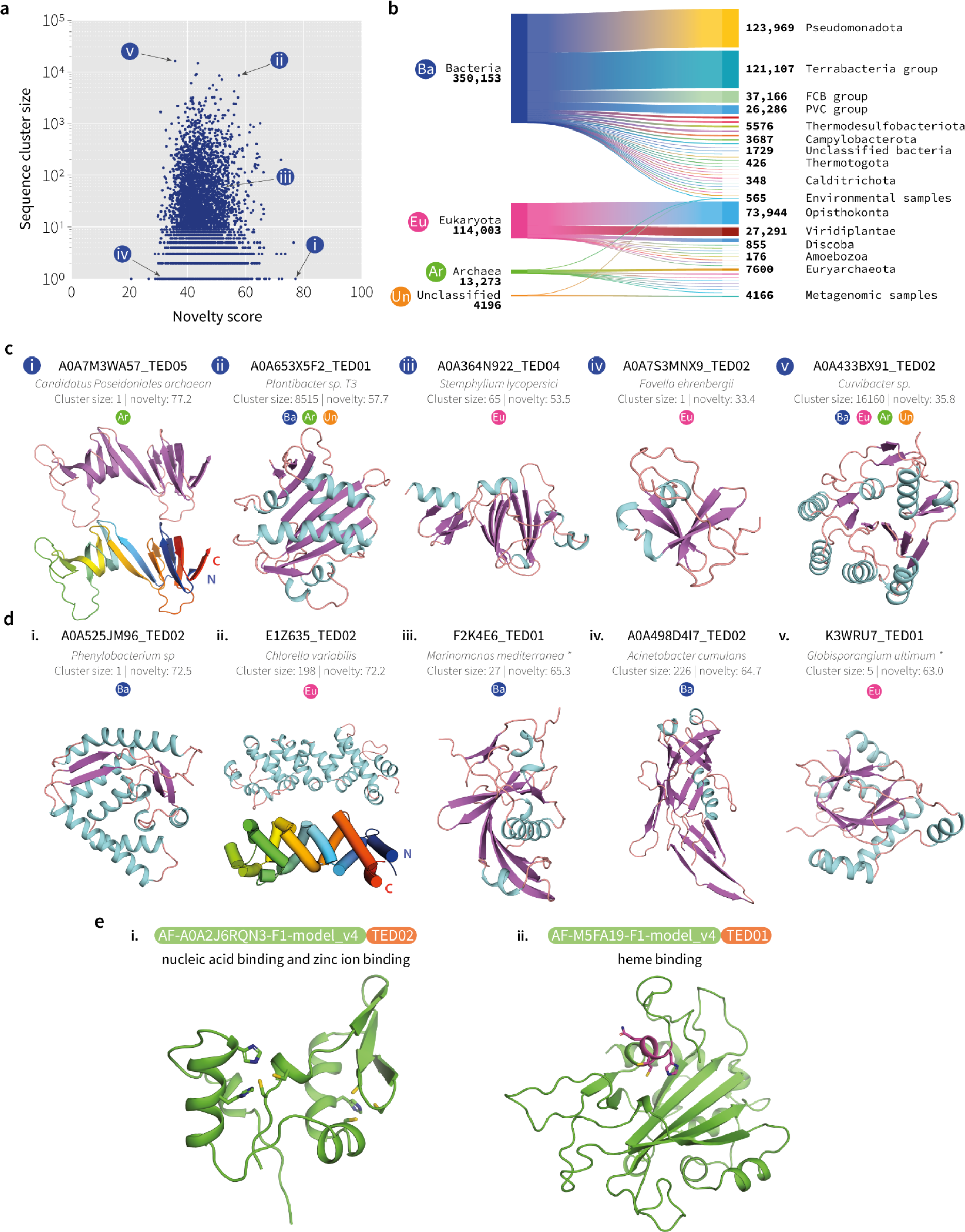
Examples of novel domain clusters identified in TED. (**a**) Comparison of domain novelty score versus sequence cluster size (n=7427). Novelty scores are predicted by the Foldclass algorithm where novel domains are ranked with a score close to 100. (**b**) Taxonomic distribution of novel domain clusters (for all sequence cluster members; n=483,732). Largest common phyla are shown across superkingdoms along with the number of domains in sequence clusters subscribed to each level of the hierarchy. (**c**) Subpanels i-v correspond to labels shown in panel (**a**). The quoted cluster size represents the number of identified homologues at the sequence cluster level. Labels denoting superkingdoms correspond to panel (**b**) and represent the superkingdom that all cluster members belong to. The cluster is distributed across multiple superkingdoms when multiple labels are shown. (**d**) Examples of high-novelty structures. In panel ii, the bottom panel shows the arrangement of helices that form the coiled hairpin loop from the N-terminus (blue) to C-terminus (red). Asterisks denote where organism names have been shortened: iii. *Marinomonas mediterranea* (strain ATCC 700492 / JCM 21426 / NBRC 103028 / MMB-1), v. *Globisporangium ultimum* (strain ATCC 200006 / CBS 805.95 / DAOM BR144) (*Pythium ultimum*). (**e**) Novel folds with predicted functions. i. Example of a domain predicted to have nucleic acid and zinc binding properties. Potential zinc binding site residues are highlighted as sticks. The left-hand site is composed of 2 Cys and 2 His residues, whereas the right-hand site has 3 Cys and 1 His in a tetrahedral arrangement. ii. Example of a heme binding domain. The residues of the heme *c* binding motif are highlighted.

Ranked first by novelty, is a curious archaeal domain found as a sequence singleton from *Candiatus Poseidoniales archaeon* (TED: A0A7M3WA57_TED05; novelty score: 77.2; Figure 4c-i). This protein is not documented in InterPro and no GO terms are available. The structure is composed of paired beta-strands, in a closed, twisted hairpin with both termini adjacent to one another. Reviewing the context of the full-chain from the AFDB shows that the domain forms part of an extended loop, protruding from the middle of a immunoglobulin-like domain (TED: A0A7M3WA57_TED04). The hairpin topology of A0A7M3WA57_TED05 is mirrored by another identified novel domain represented by E1Z635_TED02 shown in Figure 4d-ii, but is alpha-helical in nature. This domain is found only in eukaryotes, primarily in species belonging to the Viridiplantae phyla (97 species) but also in a minority of Opisthokonta (14 species) and Amoebozoa (1 species), suggesting an evolutionary link between these lineages.

### Sequence-based function prediction for novel fold and repeat domains

To see if any functions could be assigned to the domains with novel folds and repeats, we used a sequence-based deep learning model (Supp. Methods) to predict Gene Ontology (GO) terms. This analysis shows that 1321/7427 (18%) of the domains in the putative novel fold set, and 1419/6433 (22%) of the repeat set can be assigned high confidence (*p* < 10^-4^) Molecular Function GO term labels. The top 20 GO terms predicted for the two sets of domains are shown in Supp. Table 3 and 4. Manual inspection of the domains predicted to have zinc binding and nucleic acid binding functions reveals that many of the domains contain plausible zinc binding sites^28^, most containing 2 Cys and 2 His residues arranged in tetrahedral fashion, including as part of zinc finger-like supersecondary structure motifs. Figure 4e-i shows one example containing two zinc binding sites and a possible nucleic acid binding alpha-helix, but which lacks the canonical zinc finger supersecondary motif. We also consider the set of domains predicted to have heme binding properties and find that most of these contain the canonical heme *c* binding motif (CXXCH)^29^. Inspection of the three-dimensional structures reveals that each of these domains has one or more heme binding sites in a plausible conformation; one example is shown in Figure 4e-ii, with the residues of the heme *c* binding motif highlighted. The His residue that binds the heme iron is in a conformation compatible with placement of the heme group in the pocket, which is primarily hydrophobic. The presence of clear sequence motifs and structural features consistent with the assigned functions suggests that many of these novel domains may indeed have the predicted functions.

### Novel Interactions Between Domain Pairs

Unlike sequence based domain annotations such as Gene3D, the availability of full-chain AF2 models for multidomain proteins, allows us the unique opportunity to interrogate and compare domain pair packing interactions in TED and CATH. TED contains a total of 27,280,057 instances of interacting domains across 13,771 Interacting Superfamily Pairs (ISP). In contrast, the set of interacting domains in CATH consists of just 196,234 instances across 5,111 ISPs. The relative enrichment in the number of instances of ISPs common to CATH and TED is shown in Fig 6a, which shows that most of these ISPs have many more members in TED. Assessing the diversity of interaction geometry for ISPs common to TED and CATH (Methods and Figure 5b-i-ii), we find that for most ISPs, the diversity of interaction geometries in TED is consistent with that seen in CATH, indicating that on average, AF2 tends to recapitulate inter-domain geometries already seen in the PDB.

**Figure 5.**
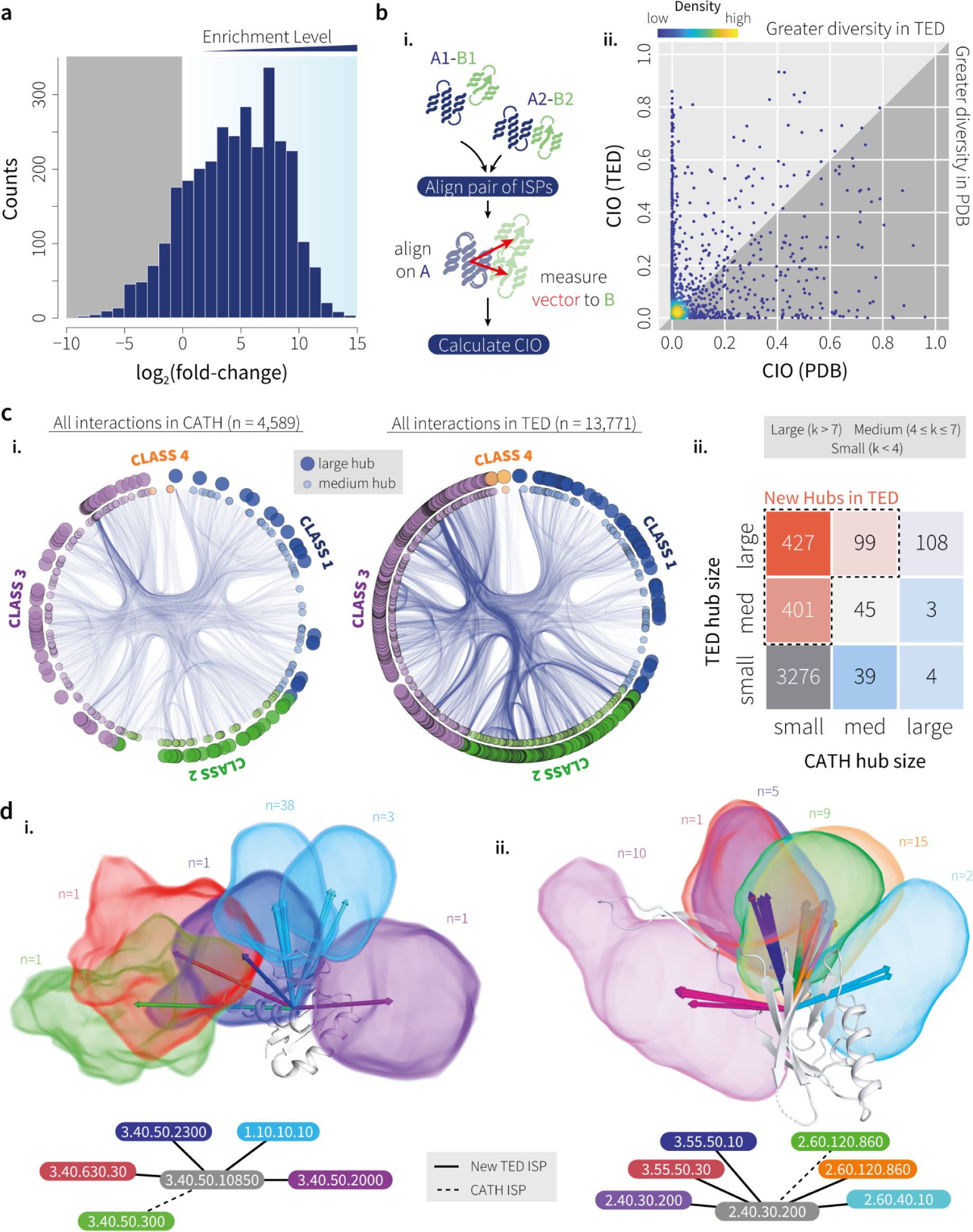
Interacting superfamily pairs (ISPs). (**a**) Enrichment of the number of instances of ISPs common to the CATH and TED datasets, expressed as log_2_(fold change) (n=3070). (**b**) i. Alignment procedure used to compute CIO values. One domain in each pair is used as reference and aligned to a designated ‘master’ reference domain. The rotation and translation from each alignment is applied to the second, ‘tag-along’ domain to bring all domain pairs into a common frame. Vectors are then computed between the centres of mass of each pair of domains, and used to compute the CIO measure (see Methods). ii. Comparison of CIO values for ISPs common to CATH and TED. Most ISPs show a high degree of conservation in interaction patterns. (**c**) i. Hierarchical edge bundling plots illustrating differences in domain superfamily interaction patterns between CATH (left) and TED (right). Curves in the plots connect interacting superfamilies. Hubs are marked by medium (4-7 connections) and large circles (>7 connections) on the outer rim. ii. Comparison of hub domains in CATH and TED. The heatmap compares CATH superfamilies in CATH and TED as hubs, categorised as small (<4 connections), medium (5-7), and large (≥8). Hub thresholds used are from Ekman et al. ^30^. (**d**) Two examples of new hub superfamilies in TED, with groups of domains for interacting superfamilies placed in a common frame and represented as volumes, alongside chains involved in each group and a graph representation of each hub. The sets of interactions for superfamily 3.40.50.10850 (NtrC-like protein domain) are disjoint between CATH and TED, whereas the set of TED interactions for superfamily 2.40.30.200 (Distal tail protein domain) include that seen in CATH (orange and green). A decomposed view of d.i. appears in Supp. Figure 14.

A small proportion (5.4%) of these ISPs show enhanced diversity in interaction geometries in TED, as measured by an increase in Conservation of Interaction Orientation (CIO) score (Methods) of 0.3 or more. A smaller proportion of ISPs (2.3%) are more diverse in CATH. Most of the interacting superfamily pairs (ISPs) in the TED set (10,701 out of 13,771) are unique to TED, i.e. they are not observed in CATH, and CATH contains 2,041 ISPs not seen in TED (Supp. Figure 13).

That all interactions in CATH are not captured in TED is not entirely surprising, firstly due to the possibility of domain parsing and classification errors, but additionally, TED excludes sequences from viruses (which are absent from the AFDB), sequences that are longer than 2700 residues, and will not contain experimental constructs comprising domain combinations not observed in natural sequences. Lastly, instances of ISPs in TED are filtered based on whether the constituent domains are in contact in the full-length AF2 structure, and whether they have a favourable inter-domain Predicted Aligned Error (PAE; Methods), both of which depend on the availability of homologous sequences and template structures that ideally span both domains. For these reasons, we do not expect that TED ISPs will be a strict superset of CATH ones. Nevertheless, we find that TED essentially doubles the set of known domain interactions at the superfamily level (10,701 novel TED interactions versus 5,111 known interactions in CATH), and further work will be needed to explore the roles that these interactions might play in cellular processes.

A visual illustration of ISP sets can be seen in Figure 5c-i, in which a path is drawn between two superfamilies if at least one interaction is observed between domains assigned to those superfamilies. This representation uses the CATH hierarchy as a guide to ‘bundle’ paths drawn between superfamilies in related parts of the CATH hierarchy (Methods; Supp. Figure 13), and shows visually that a very large number of new interactions are seen, especially between superfamilies in CATH classes 2 and 3 (all-beta and alpha-beta classes, respectively). As mentioned above, the majority of the interactions seen in the TED set are unique to TED, and a visual comparison of the set of TED-unique interactions to all known interactions in CATH (Supp. Figure 13) reveals a huge expansion in the set of known interactions. The network of superfamily interactions also allows us to identify superfamilies that can be considered hubs on account of their ability to interact with many other superfamilies. As shown in Figure 5c-i-ii, a large number of superfamilies in CATH have their hub status ‘promoted’ due to new interactions seen in TED. In Figure 5d-i-ii we show two examples of superfamilies that have been promoted in this way. Notably, the set of interactions observed in TED for these superfamilies show that they can contain only novel (in the case of 3.40.50.10850) and already-observed interaction modes (in the case of 2.40.30.200).

### Structures of redundant sequences in the AFDB

Of the 214m structures in the AFDB, nearly 39m are exact sequence duplicates of other proteins in the database (13m unique sequences within this set). These redundant proteins comprise our TED-redundant set (Methods).

Remarkably, the AFDB models for these sequence-redundant proteins often diverge from one another, with approximately 42% of clusters (5.6m) having a maximum cluster RMSD of greater than 1Å (Supp. Figure 15). The very largest RMSDs in the distribution tend to relate to changes in domain packing, but even at the domain fold level, changes can be observed. Supp. Figure 15 shows the distribution of maximum pairwise RMSD for each cluster of identical sequences. We found structural variation at the chain-level (Supp. Figure 15a), as well as in the PAE maps generated by AF2 (Supp. Figure 15a-ii).

One explanation for our observations here is that we are looking at alternative conformers of the protein chains relating to different MSAs. However, in these cases, the input MSAs should be identical given the described modelling protocol. The larger differences are also far too large to attribute to the short relaxation step that follows the model prediction step. This is most evident in the example shown in Supp. Figure 15a-ii, where two AFDB models for identical sequences deviate by nearly 65Å and show clear differences in the PAE map.

To further investigate structural diversity within these sequence-redundant clusters, we subjected the TED-redundant proteins to our domain parsing workflow, identifying approximately 69 million domains across the set. One difference employed in our treatment of the TED-redundant models is that the consensus assignment is derived across all the structures in each identical sequence cluster (Methods). This allowed us to investigate domain-level changes in conformation between identical sequences, and identified many cases where the consensus domains were dramatically different (Supp. Figure 15b).

## Discussion

What we have shown so far in developing TED is a way in which structural data in the AFDB can be augmented, by carefully breaking down structures into their component domains, allowing them to be classified through the CATH framework. This initiative not only drives forward the associations that we can make between structure and function, but as shown in our study, can be used to discover and reclaim the dark areas of fold space that are not accessible to sequence-based discovery.

Comparing TED with a recent study on the 21 model organisms dataset^6^ (released prior to the 214 million release), already shows that the TED workflow identifies not only a greater number of domains, but that the domains are also of higher quality and capture many more remote homologies (Supp. Table 2). TED currently annotates domains for over 1 million taxa, of which 600,000 are currently mapped to CATH domains in TED-100, and such an extended mapping of domains to superfamilies will enable many more evolutionary discoveries. A good example of how such expansion of CATH superfamilies enhances understanding of evolutionary processes and aids inheritance of functional properties, even towards drug repurposing is shown in Supp. Figure 16.

The coverage of TED dwarfs sequence-based assignment methods (namely Gene3D and Pfam), identifying over 100 million more domains in the AFDB compared to the latter. The proportion of proteins that each method can find domains within is shown in Supp. Figure 5, which illustrates the advantage of considering structural domains (TED finds domains in 40-50 million proteins that Gene3D and Pfam cannot). Interestingly, prior studies based purely on sequence comparisons have suggested that between 40-65% of prokaryotic proteins are composed of multiple domains, with a higher proportion proposed in eukaryotes^31–33^. These values are comparable to those seen in TED, where we find a roughly 42:55% split between single and multidomain proteins in the AFDB (Supp. Figure 5). Compared to TED, the proportion of multidomain proteins is much lower in Gene3D (29%) and Pfam (24%) assignments (Supp. Figure 5).

Although most structures in the AFDB are undeniably of high quality, the sheer scale of the data means that errors and anomalies are inevitable, and these should be discussed in order to assess the overall robustness of the data. Some limitations have already been pointed out by other studies^34,35^. One such idiosyncrasy that appeared during our development of TED was the observation that models of 100% identical sequences were sometimes dramatically different. To reframe the issue, the implication of this is that for a given protein, a user may be able to find a better model or domain within the AFDB (and one that AF2 clearly places higher confidence in) if sequence-redundant copies are considered. This may mean that some alternative structures could only be detected when redundant copies are available to compare against. Given that the vast majority of AFDB entries (175/214 million) do not have duplicates, the implication here is that an unknown (but probably large) number of low-quality structures or regions in AFDB might be improvable by resampling the input MSAs and reassessing the quality of the models. Even though this might be too heavy a task to do across the whole of AFDB, users of the database should certainly bear this in mind when looking at specific domains of interest.

The obvious explanation for the existence of divergent models for a given sequence must be that the models were generated using significantly different MSA information, and we note that explicit pre-sampling of the MSAs given to AF2 has been explored recently as a way to persuade the network to generate alternate conformations^36,37^. As the structure generation pipeline in AF2 includes possible random MSA resampling during the forward pass, it could be that a borderline MSA could result in very divergent predictions on different runs due to this resampling (fixed random seeds were not used in making the AFDB). Another possibility is that the sequence data banks used for different instances of the same target sequence were inadvertently updated to newer versions if they were predicted at later times. We were not able to investigate these possibilities further ourselves as there is no information on the MSAs provided in the AFDB data, but we do suggest that it is a topic worthy of further investigation by the AFDB developers.

Fortunately, we find many examples that reiterate the notion that plDDT is an invaluable discriminator of the confidence that a user should place on a model. Supp. Figure 17 shows an example whereby an AFDB entry has been generated within the very low confidence bin. This model features a significant number of structural defects not typical of AF2 models. “Re-folding” the sequence in ColabFold^38^ produces a number of visually striking structures across the five AF2 models, the variation of which, along with their unusually low plDDT scores, strongly indicates that AF2 has hallucinated these folds and that they should likely be disregarded.

However, overall, the large proportion of domains mapping to CATH evolutionary families gives confidence in the quality of the models, capturing well the structural features of folds and preserving their distinctive structural characteristics. In this context, the performance of the domain segmentation algorithms deployed here has also been key, as early pilot work on the 21 model organisms suggested a much higher proportion of problematic AF2 models largely caused by the poor segmentation of full-length proteins into domains by the sequence-based methods. As well as confirming earlier hypotheses that the majority of domain structural families had already been characterised experimentally^39,40^, our study has also revealed some intriguing and beautiful new domain architectures and folds, especially some highly symmetric repetitive structures.

Throughout our study, we had to make a number of algorithmic decisions which were primarily motivated by the monumental amount of data we had to process in the AFDB. As such, we intend for TED to be an ongoing development, which will evolve as the data and the needs of its users do. Our aim is to provide the community with the most comprehensive summary and breakdown of the structures within the AFDB. We expect TED to be used as a starting point for a whole host of analyses, including providing a comprehensive dataset to train and test a new generation of deep learning based applications in structural biology.

## Methods

### Datasets

Our analysis was carried out on Version 4 of the AFDB, containing models for 214,683,829 UniProt sequences. To avoid overrepresentation bias, most of our initial analysis was carried out on a non-redundant subset of 188,914,411 sequence unique AFDB model (TED-100). The 38,944,835 models which were sequence-redundant were separated into the TED-redundant dataset, with 13 million targets common between the two sets (sequence representatives were selected as the first target in a sorted list).

### Deriving a unified assignment of domains in the AFDB

Both TED-100 and TED-redundant datasets were subjected to a consensus domain parsing workflow which made use of three segmentation methods: Merizo, Chainsaw and UniDoc, before deriving a consensus between their outputs. Each method is provided with the structure of the AFDB model and returns the predicted domain ranges. In the case of UniDoc, which does not classify NDRs, models were first parsed using Merizo to remove NDRs before subsequent segmentation into domains with UniDoc. This two-step procedure for generating UniDoc domains where NDRs are first removed, is necessary as otherwise the presence of NDRs can mislead UniDoc into classifying full chains as single domains. Furthermore, removal of NDRs via using only plDDT scores is insufficient, as previously shown^20^. Overall, 400,444,974, 328,956,414 and 366,117,430 putative domain regions were identified by Merizo, Chainsaw and UniDoc respectively (Supp. Table 1 and Supp. Figure 2), requiring several months of computation on a Linux cluster. Further details on how consensus domains were derived are described in the Supp. Methods.

### Domain classification using Foldseek

Foldseek^24^ (version a435618b95ba95cbfe74dcbc8b4bd2720547d285) easy-search was benchmarked to assess tentative homology and fold assignments thresholds on a curated set of CATH domains. We created a dataset of 3,186 domains representative of CATH classes 1 to 4 clustered at 30% sequence identity with equivalent superfamily and boundary assignments in agreement with SCOP. Using a genetic algorithm, we identified thresholds for superfamily (H-level) and fold (T-level) level matches in CATH at a 98% precision level with the following parameters: E-value cutoff of 0.108662, minimum coverage of 0.366757, coverage mode 5 (shortest sequence is at least 36.6757% of the longest sequence) and sensitivity 10. Further post processing on the raw results from Foldseek was applied by using custom thresholds for H-level and T-level hits. At this precision level, the recall at H and T levels is approximately 0.59 and 0.71.

We selected as valid H-level hits any results that passed the thresholds set to Foldseek, with an additional cut-off based on a TM-score over 0.56, a coverage threshold of 0.367 and an E-value below 0.019. Hits at the fold level (T) required different cutoffs, with results considered as valid hits if passing the Foldseek E-value cutoff, a coverage cutoff of 0.786 and a minimum TM-score of 0.42. Using these thresholds, we scanned all 324,389,697 domains with a medium/high confidence against a library of 31,574 CATH SSG5 (Structural Similarity Groups, 5Å, where all CATH superfamily members are within 5Å when superposed)^9^, resulting in 193,939,494 hits at the H-level and 16,026,530 hits at the fold level, leaving 114,423,673 domains with no labels assigned at this stage.

### Domain classification using embedding similarity

To further reduce the number of unmatched domains, we subjected the remaining domains not matched by Foldseek to an in-house structure embedding search method (Merizo-search) in order to identify further matches to our CATH SSG5 representative dataset. Merizo-search uses a deep learning method called Foldclass that makes use of an equivariant graph neural network to encode a domain structure into a fixed-size embedding^25^. A query domain is embedded using Foldclass and then compared against a database of CATH domain embeddings, with similarity determined via cosine distance between the target and CATH domain embeddings. To further verify the top match, TM-align^41^ is employed to align the query coordinates against the CATH domain corresponding to the closest domain embedding. The nearest neighbour CATH domain is considered a hit if the TED domain achieves a TM-align score greater than 0.5 (normalised by the length of the TED domain).

### Sequence clustering of TED-100 domains

The sequences of TED-100 domains were clustered using MMseqs2 (version 22a77e22a77eeb1b6c64f1c5f04e1480b4705bb3fbc897), enforcing a minimum sequence identity of 50% and at least 90% coverage of the shorter sequence against the longer sequence (coverage mode 5). Sequence clusters were then populated using CATH labels identified using Foldseek and Foldclass^25^ methods (see sections ‘Domain classification using Foldseek’ and ‘Domain classification using embedding similarity’). A full breakdown of the number of labelled and unlabelled sequence clusters, and the number of domains comprising these clusters is available as Supp. Table 1.

### Novel Domain Identification Workflow

As a starting point for finding domains with putative novel folds, we filtered the remaining CATH-unlabelled clusters using three criteria to remove problematic structures and those predicted with low confidence. These filters included: two metrics that describe the globularity of the structure (radius of gyration and packing density described in Bordin et al. ^6^; Supp. Figure 18), the number of secondary structure elements, as well as a ‘plDDT_80_’ metric (Supp. Figure 19) which we make use of in our methodology. Full details on how these filters were applied can be found in the Supp. Methods.

The filtered sequence cluster representatives are then subjected to layers of further clustering and filtering (Foldseek commands are described in the Supp. Methods).

The next step of the workflow involves iterative searches of the remaining cluster representatives against the PDB and CATH^8^, ECOD^10^ and SCOPe^17^ domain libraries using Foldseek^24^. These searches use Foldseek’s easy-search using (exhaustive) TM-align mode, eliminating matches with a TM-align score greater than 0.56 and coverage of 60%. The reason for using a slightly unorthodox TM-align threshold of 0.56 is to compensate for Foldseek approximating the true TM-align score^24^, meaning that occasionally, false-positive matches are found when using the standard threshold of 0.5.

### Identifying putative novel folds

Although we consider the remaining 240,674 candidate clusters as our main working set of possible novel domain folds, further processing was carried out to try to identify a core subset of domains which were both novel and consistent with the features of well-folded protein domains. Firstly, we separated out the subset of clusters composed of high-symmetry domains. These domains may be composed of known repetitive units, for example, WD40 repeats in beta propellers, but arranged in a manner which confers novelty. High internal symmetry domains were identified using the Z-score returned by the SymD program^27^. We used a threshold of Z-score > 9.0 to indicate high-symmetry domains given the distribution shape (Supp. Figure 11).

The remaining low-symmetry clusters were assessed on domain chopping quality using a variant of the Foldclass network. This network is trained as a binary classifier to distinguish between existing CATH S30 domains and random crops of the associated full length protein chains. This classifier outputs a score between 0 to 1 which indicates the likelihood of a domain being chopped in a manner consistent with the existing CATH domains. We use a moderate threshold of quality score < 0.5 to eliminate any domains which are more likely than not of poor chopping quality, but still retain clusters that were reasonable candidates as globular domains with potentially novel folds. As an additional filter, we made use of another older domain parsing algorithm^42^ which is generally very good at identifying single domain proteins (domains that were not identified as being single domain with 100% confidence were removed).

The remaining domains were then subjected to increasingly sensitive Foldseek comparisons with the current CATH, ECOD and SCOPe domain databases^8,10,17^, with commonly-used thresholds of TM-score threshold of 0.5 and query domain coverage of 60%. The TM-score was taken as *max(qtmscore, ttmscore)* so that both the TM-scores normalised by the candidate domain length and normalised by the library domain would be considered. The final stage was to use exhaustive comparisons against the domain libraries using TM-align directly^41^. TM-align is generally more sensitive than Foldseek even in its exhaustive mode, and indeed we found further matches (TM-score > 0.5 and query coverage > 60%) to reduce the final list from 24,653 to 7,427 potential novel domain folds. This set of domains represents the set we feel most confident about defining as well-folded domains with no significant structural similarity to current PDB chains (as of December 2023) or current structural domain libraries (as of February 2024).

Using the Foldclass embeddings we ranked the final list of domains in order of novelty by calculating the mean Euclidean distance between the embedding vector of each candidate domain and the *k*-nearest neighbours (*k=50*) in CATH, ECOD and SCOPe.

### GO term analysis of novel and repeat domains

To try to evaluate the possible functions of our final repeat and novel fold domains, we used a purely sequence-based predictor (GOfocus) which makes use of a deep dilated convolutional network to predict a set of slimmed GO terms. Further details of the model can be found in the Supp. Methods.

### Evaluating domain interactions

To identify which pairs of domain superfamilies interact, and the degree of conservation in the interaction geometry for that pair of superfamilies, domains were defined as interacting if there were at least 3 Cβ atom pairs (Cα in the case of Gly residues) within 8Å between the two domains. ISPs are identified independently of the order of occurrence of the component domains in the chain sequence. For TED domains, we filter each putative set of interacting domains using the PAE data^2^ supplied with each AFDB chain prediction.

Conservation in interaction patterns was assessed using the Conservation of Interaction Orientation (CIO) measure of Littler and Hubbard^43^. A CIO score of 0 indicates complete conservation in the interaction geometry (i.e. interaction vectors for a given ISP are absolutely coincident), and a score of 1 indicates a uniformly random angular distribution of vectors. Interaction vectors were only computed for interacting domain pairs as defined above, and all instances of a given ISP are first placed in a common frame of reference by aligning on a common reference structure for one of the domains in the pair, using TM-align^41^. We also assessed the number of occurrences of ISPs common to both TED and CATH 4.3, expressing relative enrichment as the log_2_(fold change) of pairs in TED relative to CATH.

To visualise the sets of ISPs in CATH and TED, we used hierarchical edge bundling plots^44^ as implemented in the ‘ggraph’ R package^45^, using the CATH hierarchy with a virtual common root for classes 1-4 as the guide dendrogram (Supp. Figure 13). To identify hub domains, we first take each superfamily and determine the ISPs it is involved in within CATH and TED. A superfamily involved in *k* ISPs is classified as a ‘small’, ‘medium’ or ‘large’ hub when *k*<4, 4<=*k*<=7, and *k*>7, respectively, following the thresholds defined in Ekman et al.^30^ but applied to domains rather than full-length proteins.

### Validation of homology assignments using Hidden Markov Models

Consensus domain sequences with an H-level or T-level assignment by Foldseek were scanned against Hidden Markov Models built from 62,915 CATH representatives clustered at 95% sequence identity. The resulting output from HMMsearch with an E-value cut-off set at 1e-3 were subsequently post-processed using cath-resolve-hits^46^ using a minimum coverage of 80% and minimum bitscore set at 25. Resulting superfamily code assignments for H hits were checked for full matches (identical CATH codes predicted) or fold matches (first three of the CATH digits identical, different fourth level assignment between the two methods). For T-level assignments by Foldseek we compared only the first three digits in the CATH code assigned by the HMMs, returning a valid match only if the first three digits are identical, returning a non-match otherwise.

### Evaluation of domain coverage via sequence and structural searches

Full, non-redundant sequences for AFDB were scanned against Hidden Markov Models built from 62,915 CATH representatives clustered at 95% sequence identity. The resulting output from HMMsearch with an E-value cut-off set at 1e-3 were subsequently post-processed using cath-resolve-hits^46^ using a minimum coverage of 80% and minimum bitscore set at 25. The tool returns the best non-overlapping set of domain assignments on each AFDB sequence.

## Data availability

The Encyclopaedia of Domains (TED) structural domain assignments for AlphaFold Database v4 will be available as a Zenodo deposition at https://zenodo.org/records/10848710 (DOI: 10.5281/zenodo.10848710) upon publication. The deposition contains domain assignments for TED, PDB files for novel folds and individual domain assignments from Chainsaw, Merizo and UniDoc to facilitate further benchmarking efforts.

## Code availability

The code for calculating consensus domain chopping, domain quality, Foldclass embedding, search, and GO term analysis is available via the PSIPRED github repository (https://github.com/psipred/ted-tools). The code for globularity prediction is part of CATH-AlphaFlow (https://github.com/UCLOrengoGroup/cath-alphaflow). The code for the TED website is available at TED-web (https://github.com/UCLOrengoGroup/ted-web).

## Acknowledgements

We would like to acknowledge continuous support from the University College London Computer Science Technical Support Group (TSG). We also thank Sameer Velankar and Martin Steinegger for helpful discussions.

## Author Contributions

A.M.L., N.B., I.S. and D.T.J. designed and created the datasets. A.M.L. and I.S. designed and executed the domain chopping workflow. D.T.J. designed the Foldseek domain assignment process, and N.B. carried out the Foldseek domain assignment, HMM searches and sequence clustering analysis. D.T.J. designed and A.M.L. performed the Foldclass domain assignments. D.T.J. designed and executed the novel fold workflow analysis. A.M.L, D.T.J. and V.P.W. identified novel folds. A.M.L. conducted the TED-redundant analysis. S.M.K. performed the domain-domain interaction analysis. S.M.K. and D.T.J. performed GO term analysis. A.M.L. and J.W. carried out the coverage comparison analysis. D.T.J. and C.A.O. supervised the project. A.M.L., N.B., S.M.K., V.P.W., C.A.O. and D.T.J. wrote the manuscript. A.M.L, V.P.W and N.B produced figures. All authors contributed to research design and manuscript revision.

## References

1. Varadi, M. et al. AlphaFold Protein Structure Database: massively expanding the structural coverage of protein-sequence space with high-accuracy models. Nucleic Acids Res. 50, D439–D444 (2022).

2. Varadi, M. et al. AlphaFold Protein Structure Database in 2024: providing structure coverage for over 214 million protein sequences. Nucleic Acids Res. (2023) doi:10.1093/nar/gkad1011.

3. Borkakoti, N. & Thornton, J. M. AlphaFold2 protein structure prediction: Implications for drug discovery. Curr. Opin. Struct. Biol. 78, 102526 (2023).

4. Durairaj, J. et al. Uncovering new families and folds in the natural protein universe. Nature 622, 646–653 (2023).

5. Barrio-Hernandez, I. et al. Clustering predicted structures at the scale of the known protein universe. Nature 622, 637–645 (2023).

6. Bordin, N. et al. AlphaFold2 reveals commonalities and novelties in protein structure space for 21 model organisms. Commun Biol 6, 160 (2023).

7. Dustin Schaeffer, R., et al. ECOD domain classification of 48 whole proteomes from AlphaFold Structure Database using DPAM2. PLoS Comput. Biol. 20, e1011586 (2024).

8. Sillitoe, I. et al. CATH: increased structural coverage of functional space. Nucleic Acids Res. 49, D266–D273 (2021).

9. Cuff, A. L. et al. The CATH classification revisited--architectures reviewed and new ways to characterize structural divergence in superfamilies. Nucleic Acids Res. 37, D310–4 (2009).

10. Cheng, H. et al. ECOD: an evolutionary classification of protein domains. PLoS Comput. Biol. 10, e1003926 (2014).

11. Bateman, A. et al. The Pfam Protein Families Database. Nucleic Acids Res. 30, 276–280 (2002).

12. Bateman, A. et al. The Pfam protein families database. Nucleic Acids Res. 32, D138–41 (2004).

13. Lees, J. et al. Gene3D: a domain-based resource for comparative genomics, functional annotation and protein network analysis. Nucleic Acids Res. 40, D465–71 (2012).

14. Orengo, C. A. et al. CATH--a hierarchic classification of protein domain structures. Structure 5, 1093–1108 (1997).

15. Lewis, T. E. et al. Gene3D: Extensive prediction of globular domains in proteins. Nucleic Acids Res. 46, D1282 (2018).

16. Murzin, A. G., Brenner, S. E., Hubbard, T. & Chothia, C. SCOP: a structural classification of proteins database for the investigation of sequences and structures. J. Mol. Biol. 247, 536–540 (1995).

17. Fox, N. K., Brenner, S. E. & Chandonia, J.-M. SCOPe: Structural Classification of Proteins--extended, integrating SCOP and ASTRAL data and classification of new structures. Nucleic Acids Res. 42, D304–9 (2014).

18. Hadley, C. & Jones, D. T. A systematic comparison of protein structure classifications: SCOP, CATH and FSSP. Structure 7, 1099–1112 (1999).

19. Day, R., Beck, D. A. C., Armen, R. S. & Daggett, V. A consensus view of fold space: Combining SCOP, CATH, and the Dali Domain Dictionary. Protein Sci. 12, 2150–2160 (2003).

20. Lau, A. M., Kandathil, S. M. & Jones, D. T. Merizo: a rapid and accurate protein domain segmentation method using invariant point attention. Nat. Commun. 14, 8445 (2023).

21. Wells, J., Hawkins-Hooker, A., Bordin, N., Paige, B. & Orengo, C. Chainsaw: protein domain segmentation with fully convolutional neural networks. bioRxiv 2023.07.19.549732 (2023) doi:10.1101/2023.07.19.549732.

22. Zhu, K., Su, H., Peng, Z. & Yang, J. A unified approach to protein domain parsing with inter-residue distance matrix. Bioinformatics 39, (2023).

23. Steinegger, M. & Söding, J. MMseqs2 enables sensitive protein sequence searching for the analysis of massive data sets. Nat. Biotechnol. 35, 1026–1028 (2017).

24. van Kempen, M. et al. Fast and accurate protein structure search with Foldseek. Nat. Biotechnol. (2023) doi:10.1038/s41587-023-01773-0.

25. Kandathil, S. M., Lau, A. M., Buchan, D. W. A. & Jones, D. T. Foldclass and Merizo-search: embedding-based deep learning tools for protein domain segmentation, fold recognition and comparison. bioRxiv (submitted) (2024).

26. Du, D. et al. Structure of the AcrAB–TolC multidrug efflux pump. Nature 509, 512–515 (2014).

27. Tai, C.-H., Paul, R., Dukka, K. C., Shilling, J. D. & Lee, B. SymD webserver: a platform for detecting internally symmetric protein structures. Nucleic Acids Res. 42, W296–300 (2014).

28. Laitaoja, M., Valjakka, J. & Jänis, J. Zinc coordination spheres in protein structures. Inorg. Chem. 52, 10983–10991 (2013).

29. Li, T., Bonkovsky, H. L. & Guo, J.-T. Structural analysis of heme proteins: implications for design and prediction. BMC Struct. Biol. 11, 13 (2011).

30. Ekman, D., Light, S., Björklund, Å. K. & Elofsson, A. What properties characterize the hub proteins of the protein-protein interaction network of Saccharomyces cerevisiae? Genome Biol. 7, 1–13 (2006).

31. Apic, G., Gough, J. & Teichmann, S. A. Domain combinations in archaeal, eubacterial and eukaryotic proteomes. J. Mol. Biol. 310, 311–325 (2001).

32. Batey, S., Nickson, A. A. & Clarke, J. Studying the folding of multidomain proteins. HFSP J. 2, 365–377 (2008).

33. Zhou, X., Hu, J., Zhang, C., Zhang, G. & Zhang, Y. Assembling multidomain protein structures through analogous global structural alignments. Proc. Natl. Acad. Sci. U. S. A. 116, 15930–15938 (2019).

34. Hou, Y., Xie, T., He, L., Tao, L. & Huang, J. Topological links in predicted protein complex structures reveal limitations of AlphaFold. Commun Biol 6, 1098 (2023).

35. Jones, D. T. & Thornton, J. M. The impact of AlphaFold2 one year on. Nat. Methods 19, 15–20 (2022).

36. Wayment-Steele, H. K. et al. Predicting multiple conformations via sequence clustering and AlphaFold2. Nature 625, 832–839 (2024).

37. Del Alamo, D., Sala, D., Mchaourab, H. S. & Meiler, J. Sampling alternative conformational states of transporters and receptors with AlphaFold2. Elife 11, (2022).

38. Mirdita, M. et al. ColabFold: making protein folding accessible to all. Nat. Methods 19, 679–682 (2022).

39. Bordin, N., Sillitoe, I., Lees, J. G. & Orengo, C. Tracing Evolution Through Protein Structures: Nature Captured in a Few Thousand Folds. Front. Mol. Biosci. 8, 668184 (2021).

40. Chothia, C. One thousand families for the molecular biologist. Nature Publishing Group UK http://dx.doi.org/10.1038/357543a0(1992) doi:10.1038/357543a0.

41. Zhang, Y. & Skolnick, J. TM-align: a protein structure alignment algorithm based on the TM-score. Nucleic Acids Res. 33, 2302–2309 (2005).

42. Taylor, W. R. Protein structural domain identification. Protein Eng. 12, 203–216 (1999).

43. Conservation of Orientation and Sequence in Protein Domain–Domain Interactions. J. Mol. Biol. 345, 1265–1279 (2005).

44. Holten, D. Hierarchical edge bundles: visualization of adjacency relations in hierarchical data. IEEE Trans. Vis. Comput. Graph. 12, 741–748 (2006).

45. Pedersen, T. L. ggraph: An Implementation of Grammar of Graphics for Graphs and Networks. Preprint at https://ggraph.data-imaginist.com (2024).

46. Lewis, T. E., Sillitoe, I. & Lees, J. G. cath-resolve-hits: a new tool that resolves domain matches suspiciously quickly. Bioinformatics 35, 1766–1767 (2018).

