## Supplementary Document for "Exploring structural diversity across the protein universe with The Encyclopedia of Domains"

### Supplementary Materials and Methods

Lau, A. M.[1 † +], Bordin, N.[2 †], Kandathil, S. M.[1], Sillitoe, I.[2], Waman, V. P.[2], Wells, J.[2,3], Orengo, C.[2*] and Jones, D. T.[1,2*]

† These authors contributed equally to this work

**Affiliations**
[1]Department of Computer Science, University College London, London, WC1E 6BT, UK
[2]Institute of Structural and Molecular Biology, University College London, London, WC1E 6BT, UK
[3]Centre for Artificial Intelligence, University College London, London, WC1V 6BH, UK

**Present address**
[+]InstaDeep Ltd, 5 Merchant Square, London, W2 1AY, UK

### Supplementary Methods

#### Deriving a unified assignment of domains in the AFDB

The predicted domains on full length AFDB targets by Merizo, Chainsaw and UniDoc were filtered to remove any segments that are fewer than 5 residues, and any domains which are fewer than 25 residues. To determine a common set of putative domains identified by all three methods, domains are classified into high, medium or low consensus categories depending on the level of agreement between the predictions of the three methods (Supp. Figure 3). Specifically, given a set of N domain predictions for a given target, we construct an N-by-N matrix of pairwise sequence overlap scores between each domain, thresholded at 70% coverage to generate an adjacency matrix. This matrix can be treated as a network graph, with the size of each internal graph component indicative of the confidence level (3, 2 and 1 indicating high, medium and low confidence, respectively).

For TED-100, of which there are 188m unique sequences, the consensus is derived for each individual protein. High, medium and low confidence categories reflect the agreement of predictions by either three, two or single methods. The final domain ranges reported, are taken as the intersection between the regions in agreement with each other. For domain counts and subsequent downstream analysis tasks, only the high and medium categories are considered due to their non-overlapping nature. Due to the nature of their derivation, high and medium consensus domains cannot overlap.

For TED-redundant, where each unique sequence is associated with a variable number of models, the consensus is derived across predictions made on all constituent models. This also necessitates dynamic thresholding for high and medium consensus categories due to the variable number of models being considered for each sequence. Thresholds were derived as *threshold = 3Mk*, where *M* is the number of targets (2 minimum for TED-redundant), and *k* is either 0.66 or 0.33 for high and medium confidence levels, respectively. For the TED-redundant set only, high consensus domains are given priority over those of medium consensus, with any in the latter category overlapping with high consensus domains disqualified.

Using this approach, we identified 324,482,131 domains in TED-100 (195,229,271 at high consensus and 129,160,426 at medium consensus) and 68,764,894 domains in TED-redundant (58,002,177 at high consensus and 10,762,717 at medium consensus). As 13m targets are shared across TED-100 and TED-redundant, the final adjusted domain count across the AFDB is 371,357,492 domains.

**Novel Domain Identification Workflow**

Our novel domain identification workflow makes use of several filters that were applied to non-CATH-labelled cluster representatives. First, the representatives of the 41,879,858 domain clusters were assessed using the normalised radius of gyration and packing density described in Bordin et al.[1]**.** Globular TED domains were determined with CATH-AlphaFlow as those satisfying both criteria of having a normalised radius of gyration of below 0.356, and a packing density of greater than 10.333 (Supp. Figure 18). Cutoff values were determined as the 5th percentile of both metrics calculated from the 193,939,492 TED-100 domains with H-level CATH assignments, as these domains were of sufficient quality to be representative of domains in the AFDB.

For secondary structure elements (SSE) filtering, domains were excluded from consideration as novel domains if they contained fewer than 6 SSEs (helices and strands), as determined by STRIDE[2,3]. STRIDE counts even very short regions (2 residues) as separate secondary structures, so a threshold of 6 elements in total was deemed a sensible threshold to allow us to identify novel domain architectures. Applying both the globularity and SSE filters to the 41,879,858 unlabelled clusters, results in 13,820,550 passing both criteria.

Finally, we apply a plDDT filter on the remaining domains, which we term the ‘$\text{plDDT}_{80}$’ metric. The $\text{plDDT}_{80}$ is calculated as the average plDDT of the top 80% of residues ranked by plDDT. We found $\text{plDDT}_{80}$ preferable to the simple average as it prevented removing clusters that are mainly well-folded but have unfolded tail regions. Applying a $\text{plDDT}_{80}$ >= 90 filter on the remaining 13,820,550 clusters yields a final set of 8,612,318 clusters which pass all three (globularity, SSE and $\text{plDDT}_{80}$) criteria, encompassing 19,816,697 domains within them.

**GO term analysis of novel and repeat domains**

To try to evaluate the possible functions of our final repeat and novel fold domains, we used a purely sequence-based predictor called GOfocus, which performed well in the 4th CAFA experiment[4] [https://biofunctionprediction.org/cafa/]. The model comprises a 1-D dilated convolutional network with 4 blocks of 4 dilated convolutional layers per block, with dilations cycling through 1, 2, 4 and 8, a filter size of 3, 512 channels and residual connections across each layer. After the dilated convolutional blocks, adaptive max-pooling is used to reduce the (L,512) tensor to fixed dimensions (16,512) which are then flattened and used as input features for a two layer fully connected network with ReLU non-linearity and dropout rate set to 0.1. Sixteen separate networks were trained with different non-overlapping subsets of a slimmed set of 8360 GO term labels. Subsets were chosen to keep terms with similar observed frequencies in the sequence data bank grouped together. Training was carried out on the UniRef50 subset of sequences in the December 2019 release of SwissProt and TrEMBL and their respective (non-IEA) GO terms[5]. Focal loss was used as the loss function and training was carried out with the standard Adam optimizer with a learning rate of 1e-4. Testing this model on the CAFA3 benchmark test data produced an $F1_{max}$ score of 0.65 for Molecular Function terms.

**MMseqs commands for sequence clustering**

Number of sequences input to clustering: 324,389,698
Number of clusters generated: 120,748,700

Clusters generated using:

```
mmseqs easy-linclust domain_sequences.fasta results tmp --cov-mode 5 -c 0.9
--min-seq-id 0.5
```

**Foldseek commands used for novel fold identification workflow**

Number of models input to clustering: 8,612,318
Number of clusters generated: 1,036,525

Clusters generated using:

```
foldseek cluster foldseek.db result tmp -s 4 -e 0.001 -c 0.7 --cov-mode 5
--tmscore-threshold 0.56 --min-seq-id 0.2 --seq-id-mode 1 --cluster-mode 1
```

1,036,525 clusters were further clustered using below command to generate 625,802 clusters:

```
foldseek easy-cluster models result tmp -c 0.8 --tmscore-threshold 0.8
--cluster-mode 1
```

625,802 cluster representatives searched against PDB chains using:

```
foldseek easy-search models PDB result.m8 tmp -c 0.6 --cov-mode 2
--tmscore-threshold 0.5 --alignment-type 1 --format-output
query,target,qlen,tlen,qtmscore,cigar
```

Output file filtered using `qtmscore > 0.56` and cigar-computed `qcov > 0.6` to generate resultant 358,491 clusters. Clusters are searched against domain libraries using:

```
foldseek easy-search models domain_db result.m8 /ssd1/jones/tmp -c 0.6
--tmscore-threshold 0.3 --alignment-type 1 --exhaustive-search 1 --format-output
query,target,qlen,tlen,qtmscore,cigar
```

Results are filtered using `qtmscore > 0.56` and cigar-computed `qcov > 0.6` and cigar-computed `tcov > 0.6` to generate 240,674 clusters.

Low symmetry domains comprising 138,120 clusters (after domain quality and high-symmetry filtering) are clustered together with domain libraries:

```
foldseek easy-cluster models_with_domain_db result tmp --tmscore-threshold 0.5
-c 0.6 --alignment-type 1
```

This generates 39,037 clusters which are not composed of any members from the domain databases.

39,037 cluster representatives are searched against domain libraries using:

```
foldseek easy-search models domain_db result.m8 tmp -c 0.6 --cov-mode 2
--tmscore-threshold 0.3 --alignment-type 1 --exhaustive-search 1 --format-output
query,target,qlen,tlen,qtmscore,ttmscore,rmsd,cigar
```

Results are filtered for `max(qtmscore,ttmscore) > 0.5` or `RMSD < 3` and cigar-computed `qcov > 0.6`, retaining 24,653 clusters.

Final clusters are searched again against domain libraries using manual TM-align runs and filtered using `qcov > 0.6` and `max(qtmscore,ttmscore) > 0.5` to leave 7,427 final clusters.

**Detection of poor quality domain choppings using an EGNN network**

Although the use of a consensus domain segmentation method produces generally good quality domains, inevitably, amongst the long tail of possible novel folds we will find candidate domains that represent either over- or under-chopped chain segments. By under-chopped we mean structures that still appear to include domain linker regions, and by over-chopped we mean structures that are similar to existing domains but which are incomplete e.g. a segment of a TIM-barrel. As such, we assessed each remaining low-symmetry cluster on chopping quality, by retraining a variant of the Foldclass network to identify tenuous domain choppings.

We employed a version of the Foldclass network (with 3 EGNN layers rather than 2) to score domains in terms of the quality of the segmentation with a two class (binary) output layer.

For every domain in the 30% non-redundant CATH 4.3 domain set, we trained the network to recognize these domains as being in the positive class i.e. are correctly segmented. For the negative labelled cases, we generated an entirely random crop of the related full-length protein chain. A random 50-50 mix of positive and negative labelled cases was considered during each epoch of training.

#### Structural characterisation of CoA-dependent acyltransferases in TED

CATH domains for the Chloramphenicol Acetyltransferase superfamily (3.30.559.10, n=254) were scanned in an all-vs-all fashion using SSAP[6] and clustered with complete-linkage using cath-cluster (https://cath-tools.readthedocs.io/en/latest/tools/cath-cluster/) into 14 SSG5 (Structural Similarity Groups, 5Å), which were inspected for oligomerization state and ligand data available in the Protein Data Bank. TED domain structures with an associated H-level assignment to the 3.30.559.10 superfamily were retrieved (n=228,867) and scanned against the library of 14 SSG5s using Foldseek-TMalign with the following command:

```
foldseek easy-search pdb/ 3.30.559.10_ted_domains_models.tar.gz ssg5_vs_ted_domains_tmalign_over_0.7 tmp --max-seqs 300000 --cov-mode 5 -c 0.6 --tmscore-threshold 0.7 --alignment-type 1 --tmalign-hit-order 1 --format-output query,target,qstart,qend,qlen,qcov,tstart,tend,tlen,tcov,alnlen,pident,qtmscore,ttmscore,rmsd
```

TED hits retrieved from Foldseek were annotated with a comprehensive set of pathogens-associated TaxonIDs (n=39,664) from BV-BSRC v3.35.5 (https://www.bv-brc.org/), ENA Pathogens Portal (https://www.pathogensportal.org/) and VEuPathDB (https://veupathdb.org/veupathdb/app/). Hits were subsequently sorted by sequence similarity, TMscore, pathogenicity and pIDDT, with priority for analysis given to SSGs with known ligand information in PDB, low sequence similarity to any known PDB structures, high TMscore for both query and target, and high pIDDT.

For CATH experimental domains, known substrate binding residues were retrieved from the literature and PDBSum[7]. For TED domains, predicted conserved residues were obtained by scanning the TED domain against AlphaFold/UniProt50 v4 using Foldseek server (https://search.foldseek.com/search), retrieving structural relatives with TM-Score greater than 0.70 and where conserved sites were detected by the Scorecons algorithm[8] on the multiple sequence alignment derived from the relatives.

The trimer for the TED domain in *Clostridium botulinum* was modelled with AlphaFold-Multimer using ColabFold 1.5.2[9] with 20 recycles and amber relaxation using the following command:

```
colabfold_batch --templates --amber --num-models 5 --num-relax 5
```

### Supplementary Tables

#### Supplementary Table 1. Breakdown of TED workflow.

| **Step** | **No. domains** | **No. Clusters** |
|---|---|---|
| **i. Datasets** | | |
| No. targets (full AFDB) | 214,683,829 | 188,914,411 |
| TED-100 | 188,914,411 | |
| TED-redundant | 38,944,835 | |
| **ii. Domain assignment (TED-100)** | | |
| Raw domains (Merizo) | 400,444,974 | |
| Raw domains (Chainsaw) | 328,956,414 | |
| Raw domains (UniDoc) | 366,117,430 | |
| Medium consensus | 129,160,426 | |
| High consensus | 250,629,037 | |
| Total no. domains (TED-100) | 324,389,697 | |
| **iii. Classification** | | |
| Superfamily labels assigned | 193,939,494 | |
| Topology labels assigned | 45,796,122 | |
| Domains with no labels | 84,654,081 | |
| Sequence clusters (50% identity, 90% coverage) | 324,389,697 | 120,748,700 |
| Non-singletons | 242,723,300 | 39,082,303 |
| with CATH label | 202,318,609 | 29,872,777 |
| without CATH label | 40,404,691 | 9,209,526 |
| Singletons | 81,666,397 | 81,666,397 |
| with CATH label | 48,996,065 | 48,996,065 |
| without CATH label | 32,670,332 | 32,670,332 |

| | | |
|---|---|---|
| Total CATH assignable | 251,314,674 | 78,868,842 |
| Annotations transferrable | 11,579,058 | |
| Total unassigned | 73,075,023 | 41,879,858 |
| Passing globularity, nSSE, plDDT filters | 19,816,697 | 8,612,318 |
| Discarded by filters | 53,258,326 | 33,267,540 |
| **iv. Novel fold workflow** | | |
| Structures considered | 19,816,697 | 8,612,318 |
| Structure clusters | 8,612,318 | 625,802 |
| Clusters matched to PDB, ECOD, SCOPe and CATH | 13,297,059 | 385,128 |
| Unmatched clusters | 6,519,638 | 240,674 |
| High internal symmetry clusters | 277,694 | 6,433 |
| Putative novel clusters (low symmetry) | 3,614,884 | 138,120 |
| Final novel clusters (after final searches) | 483,732 | 7,427 |

**Supplementary Table 2. Comparison of coverage between Bordin et al., 2023 and TED for AFDB 21 model organisms dataset.**

| Source | AF21 | AF21 (%) | TED21 | TED21 (%) |
|---|---|---|---|---|
| Chopped domains | 708,941 | | 1,300,686 | |
| Domains over 70 plDDT | 532,412 | 75.04% | 1,176,636 | 90,46% |
| Good quality domains | 369,512 | 52.08% | 867,174 | 66,67% |
| Good quality domains in CATH | 341,213 | 92.3% | 718,703 | 82.87% |

The characterisation of the initial release of the AFDB, described in Bordin et al.[1], increased the structural coverage for 21 model organisms with over 341,000 domains with good quality assigned to CATH. Using the same thresholds specified in the original article on TED domains associated to proteomes from the original release, we noticed how TED nearly doubles the number of chopped domains, and its consensus approach identifies a higher proportion of domains with high quality (AF21=52%, TED21=67%). The percentage of good quality domains assigned to CATH with TED is slightly lower whilst the total count is higher than AF21, suggesting that the original AF21 dataset comprised mostly close homologs, while TED encompasses those and further globular domains with more remote relationships.

**Supplementary Table 3. Top 20 most frequent GO molecular function terms predicted in the set of domains with novel folds.**

| GO term | Count | Description |
|---|---|---|
| GO:0016491 | 386 | oxidoreductase activity |
| GO:0016301 | 120 | kinase activity |
| GO:0008270 | 110 | zinc ion binding |
| GO:0008168 | 101 | methyltransferase activity |
| GO:0003676 | 87 | nucleic acid binding |
| GO:0008233 | 80 | peptidase activity |
| GO:0020037 | 72 | heme binding |
| GO:0016874 | 53 | ligase activity |
| GO:0004519 | 43 | endonuclease activity |
| GO:0016829 | 39 | lyase activity |
| GO:0008237 | 25 | metallopeptidase activity |
| GO:0009055 | 25 | electron transfer activity |
| GO:0004222 | 23 | metalloendopeptidase activity |
| GO:0016757 | 23 | glycosyltransferase activity |
| GO:0051536 | 22 | iron-sulphur cluster binding |
| GO:0005509 | 21 | calcium ion binding |
| GO:0022857 | 21 | transmembrane transporter activity |
| GO:0003723 | 20 | RNA binding |
| GO:0003968 | 19 | RNA-dependent RNA polymerase activity |

**Supplementary Table 4. Top 20 most frequent GO molecular function terms predicted in the set of novel repeat domains.**

| GO term | Count | Description |
|---|---|---|
| GO:0016491 | 278 | oxidoreductase activity |
| GO:0016301 | 127 | kinase activity |
| GO:0022857 | 89 | transmembrane transporter activity |
| GO:0008270 | 87 | zinc ion binding |
| GO:0003677 | 79 | DNA binding |
| GO:0003700 | 77 | DNA-binding transcription factor activity |
| GO:0003676 | 76 | nucleic acid binding |
| GO:0016829 | 74 | lyase activity |
| GO:0008233 | 59 | peptidase activity |
| GO:0008168 | 56 | methyltransferase activity |
| GO:0005509 | 47 | calcium ion binding |
| GO:0016874 | 43 | ligase activity |
| GO:0004650 | 36 | polygalacturonase activity |
| GO:0020037 | 36 | heme binding |
| GO:0043565 | 34 | sequence-specific DNA binding |
| GO:0016757 | 27 | glycosyltransferase activity |
| GO:0030246 | 26 | carbohydrate binding |
| GO:0009055 | 25 | electron transfer activity |
| GO:0016853 | 24 | isomerase activity |
| GO:0046872 | 19 | metal ion binding |

### Supplementary Figures

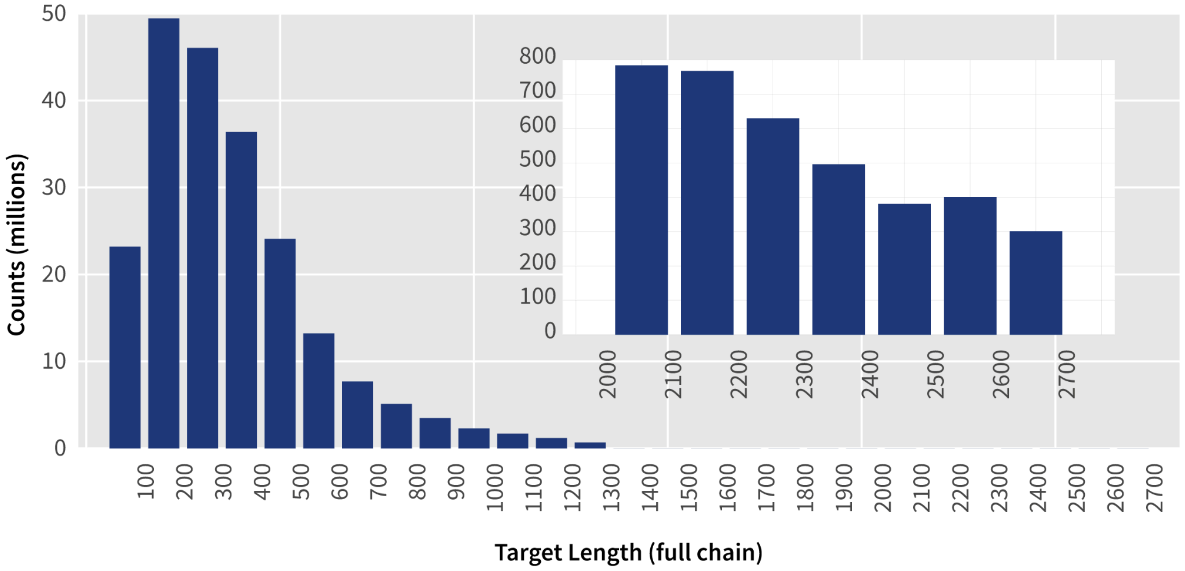

**Supplementary Figure 1. Distribution of target lengths in the AFDB.** Histogram encompasses the full database of full length targets in the AFDB (n=214,683,829). Inset shows the distribution of target lengths for targets longer than 2000 residues. The AFDB contains models up to a maximum length of 2700 residues (as part of the full distribution containing 214m targets).

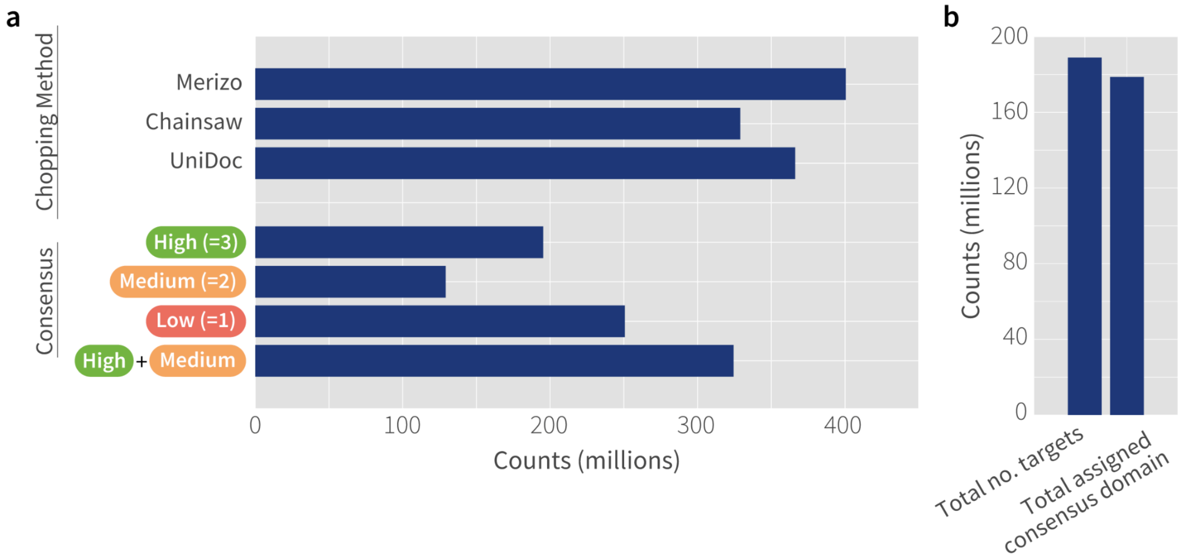

**Supplementary Figure 2. Domain counts identified from consensus chopping methods in TED-100.** (a) Number of domains identified from Merizo, Chainsaw and UniDoc methods. Domain regions were aggregated into three consensus levels: high (agreement between all three methods), medium (agreement between two methods) and low (regions without agreement between at least two methods). Medium and high consensus domains were taken forward for analysis. (b) Number of targets in TED-100 with consensus domain (high and medium levels) coverage. 98% of targets are covered by TED-100 in total.

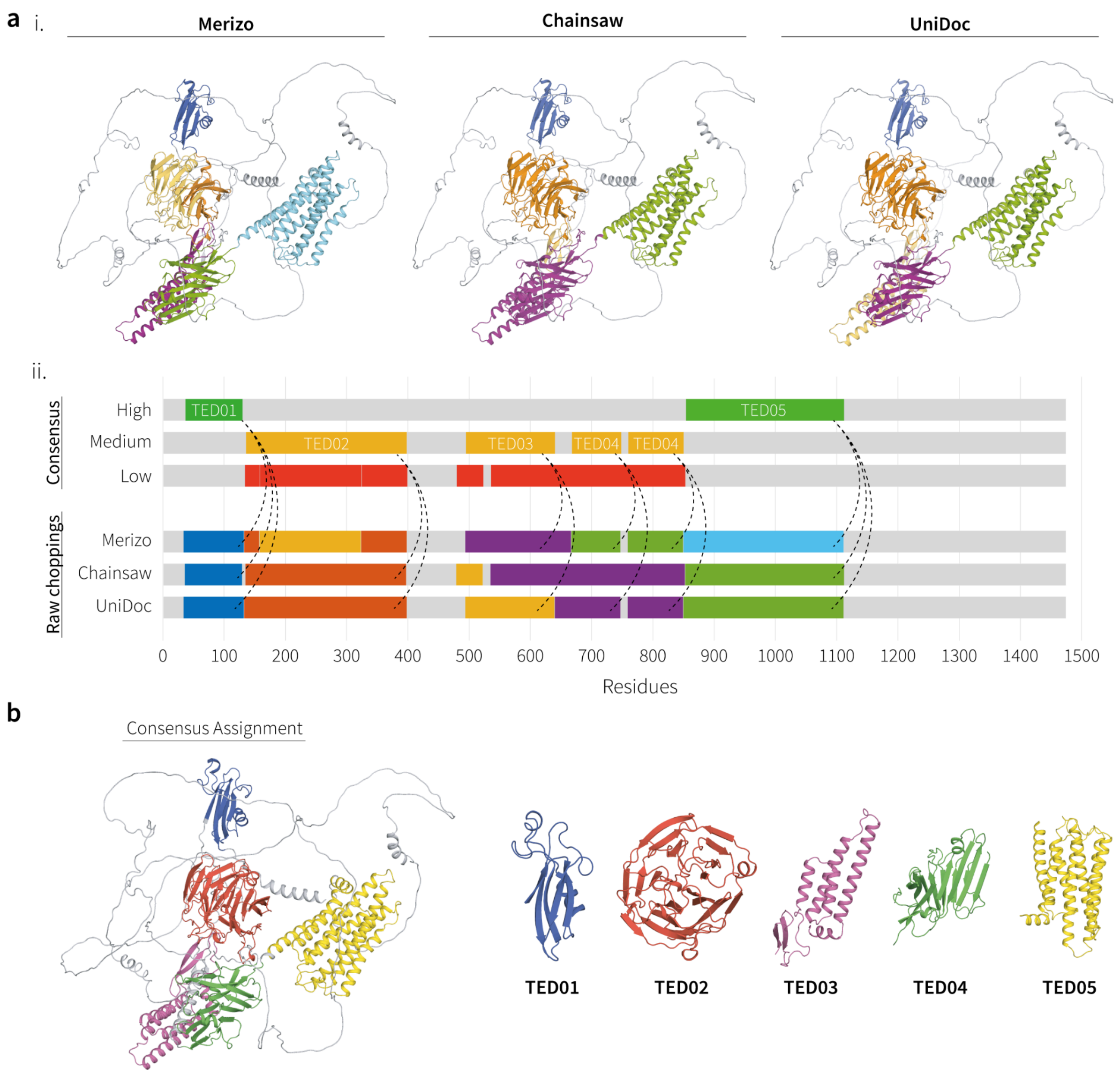

**Supplementary Figure 3. Example of consensus domain derivation.** (**a**) i. Example of predicted domain choppings from Merizo, Chainsaw and UniDoc for target AF-O94910-F1-model_v4 (Adhesion G protein-coupled receptor L1 from *Homo sapiens*). Predicted domains are shown in alternating colours. ii. Domain ranges in panel i. shown in one dimension along with the consensus classification for predictions across the three methods. High and medium consensus domains are those where three and two predictions from constituent methods agree. Low consensus domains are any single-method only predictions with low support and are not part of TED-100. Agreement between domain ranges are calculated based on a minimum of 70% sequence overlap between multiple domain ranges. Dashed lines have been drawn on the consensus panel to indicate source ranges used to derive the consensus from for medium and high confidence categories. TED domains are named sequentially beginning from the N-terminus. (**b**) Example of the consensus assignment for target AF-O94910-F1-model_v4 derived in panel (a), alongside views of individual identified domains. Domains are shown in alternating colours.

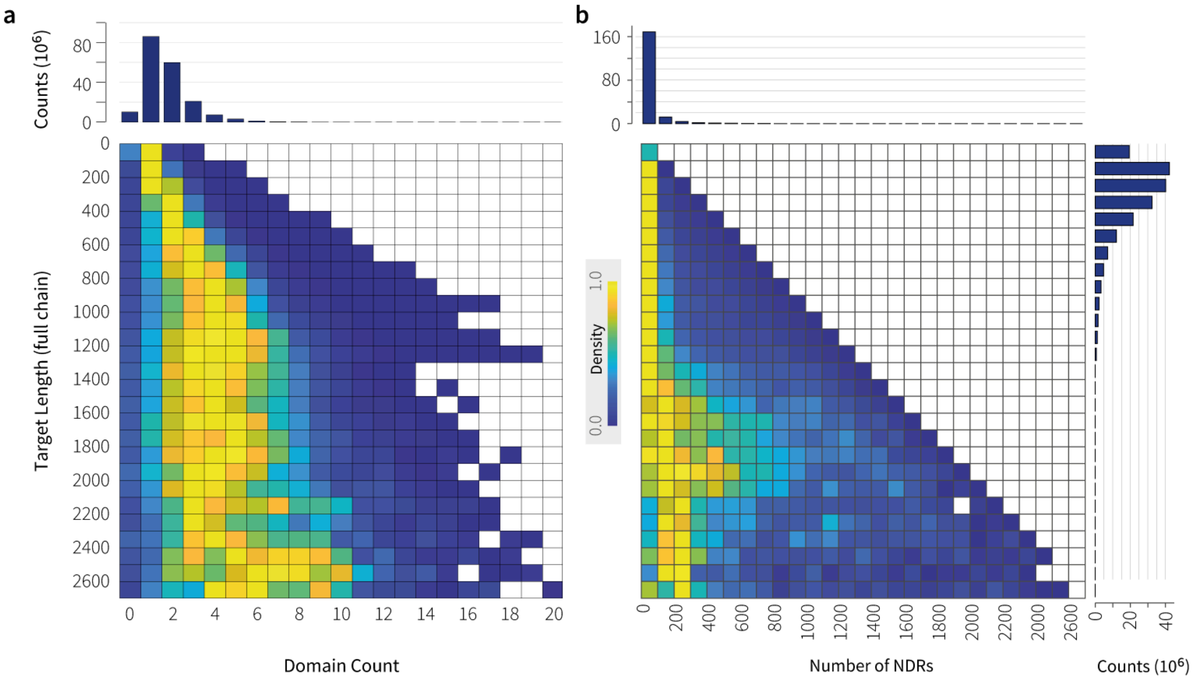

**Supplementary Figure 4. Distribution of domain counts and number of NDRs across targets in TED-100.** Data encapsulates 188m full length AFDB targets in TED-100. (**a**) Domain counts per target were taken as the total number of medium and high consensus domains. (**b**) NDRs were determined as residues not assigned into any domains by either of the three chopping methods (residues not in any low, medium or high consensus categories). Colour scheme has been normalised across rows. Each row represents a bin of 100 residues, up to the maximum length of 2700 for AFDB models.

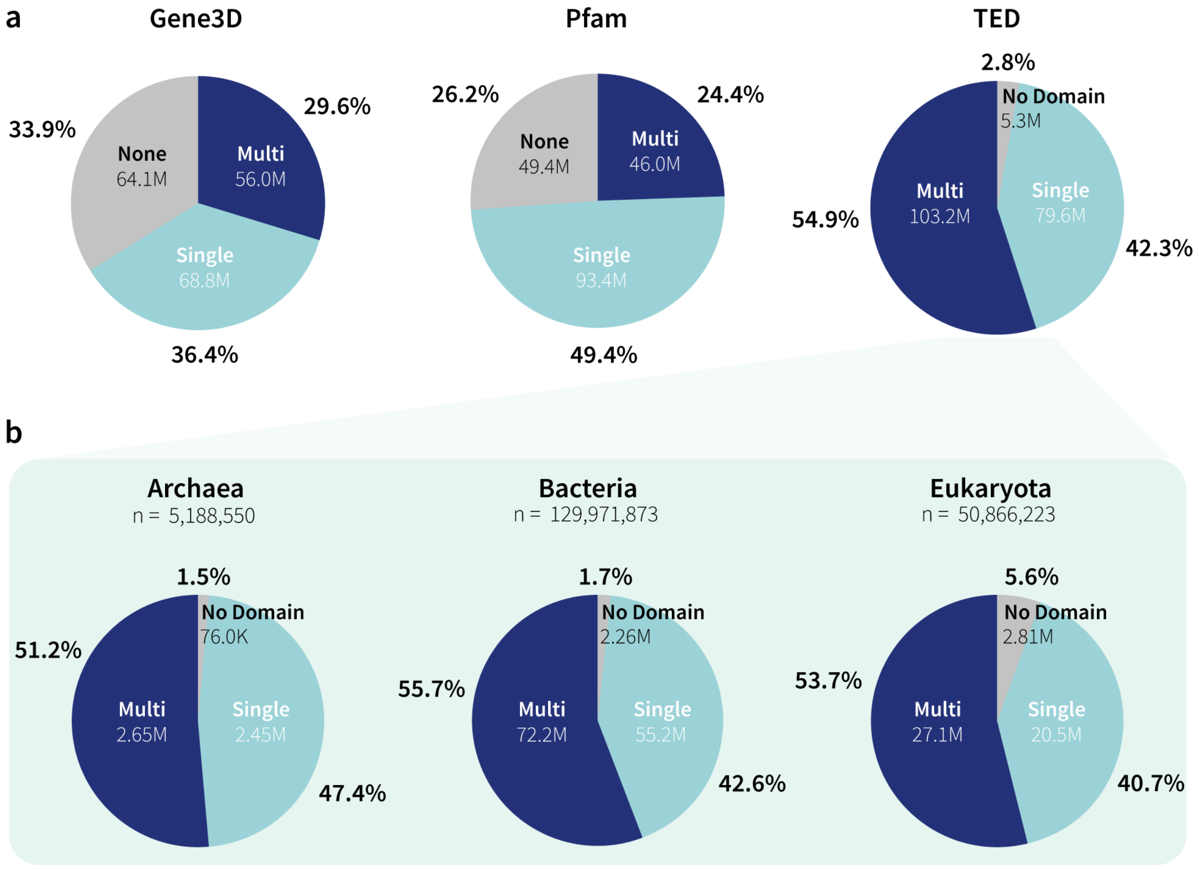

**Supplementary Figure 5. Proportion of single- and multi-domain targets identified in TED-100.** (**a**) Comparison of domain compositions according to Gene3D, Pfam and TED assignments. (**b**) TED-100 assignments subdivided by the three major superkingdoms.

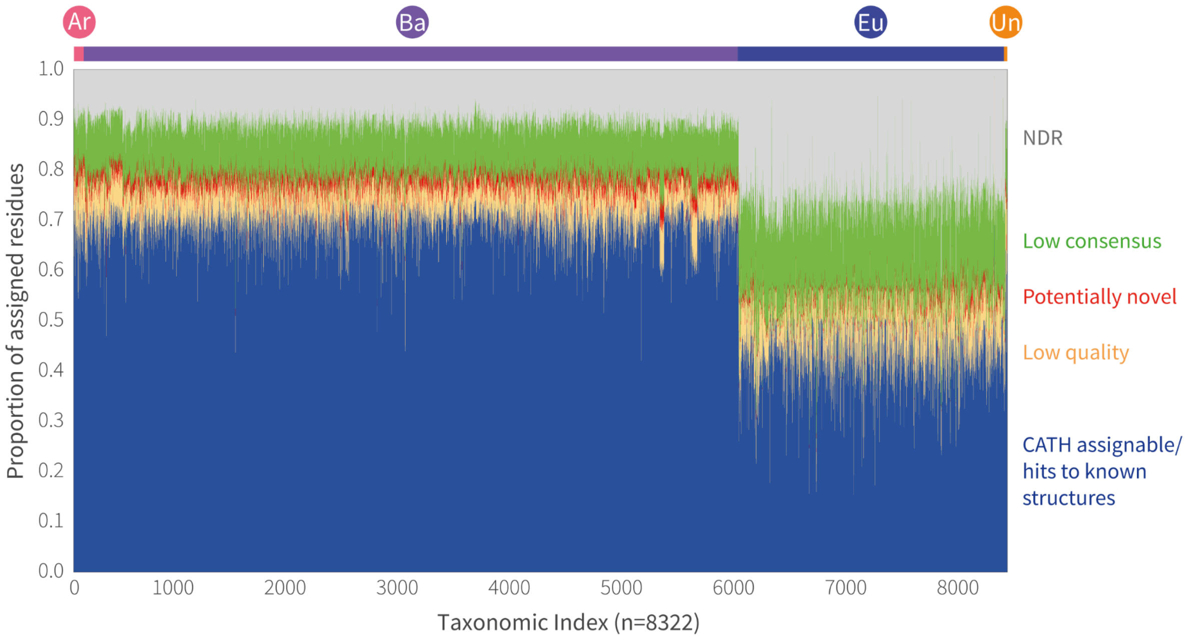

**Supplementary Figure 6. Proportion of assigned residues in TED-100.** Data shown represents a subset of taxa which are composed of at least 5000 targets across TED-100 (n=8322). Taxa are grouped based on UniProt superkingdoms (archaea (Ar), bacteria (Ba), eukaryota (Eu) and unclassified (Un)). “CATH assignable” (blue) includes domains from sequence clusters containing at least one CATH-assigned member, as well as domains that can be matched to PDB and domain databases. “Low quality” (yellow) encapsulates domains removed due to low-globularity, few-SSE or low-plDDT filters. “Novel” (red) category includes domains considered as novel domains. “Low consensus” (green) category includes residues where medium/high consensus could not be reached, but are non-NDR. “NDR” (grey) represents residues not considered to be part of domains by any of the three domain segmentation methods.

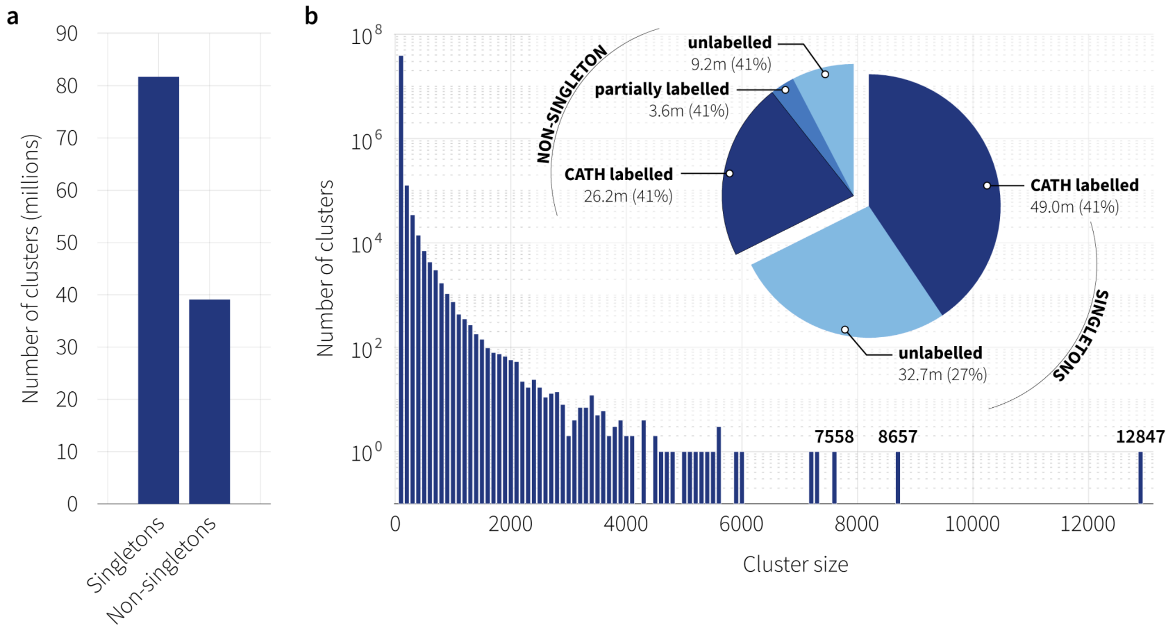

**Supplementary Figure 7. Summary of sequence clusters in TED-100.** (**a**) Number of singleton and non-singleton domain sequence clusters identified at 50% sequence identity and minimum overlap of 90%. (**b**) Histogram of cluster sizes for non-singleton clusters (bin width of 100). The majority of clusters are encapsulated within the first bin (below 100 members). The number of sequence members in the largest 3 bins are displayed. Inset shows the proportion of singleton and non-singleton clusters that can be assigned CATH labels. 'CATH labelled' clusters represent clusters where every member is assigned a CATH label at the superfamily or topology levels. 'Partially labelled' clusters contain at least one member with a CATH superfamily or topology level. 'Unlabelled' clusters are entirely unlabelled.

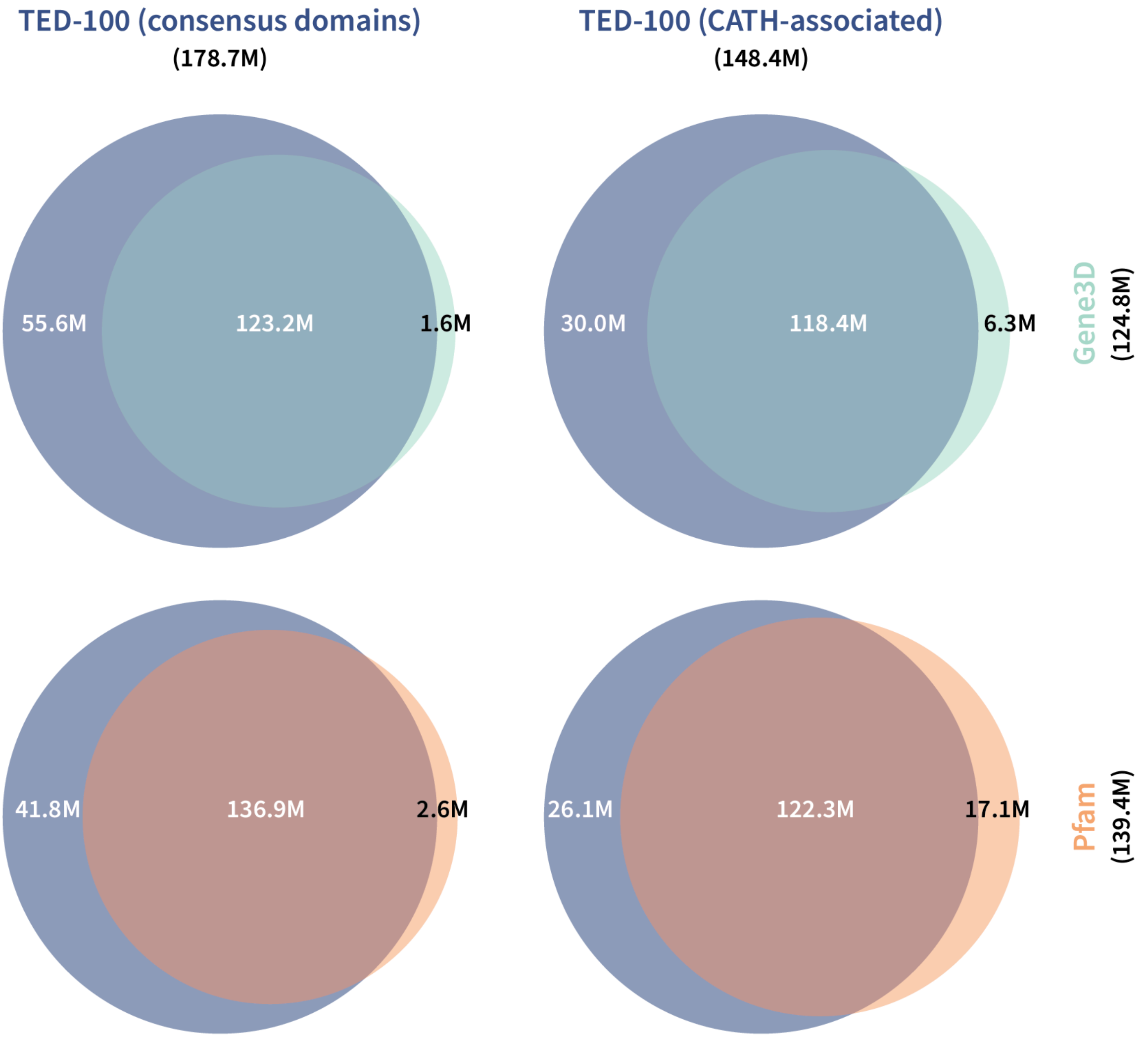

**Supplementary Figure 8. Comparing the coverage of TED against Pfam and Gene3d.** Venn diagrams compare the protein-level coverage of TED-100, across all proteins encompassing high and medium consensus domains (178.7 million proteins), and those that are associated with CATH-labelled clusters (148.4 million proteins), with those of Gene3D (124.8 million proteins) and Pfam (139.4 million proteins).

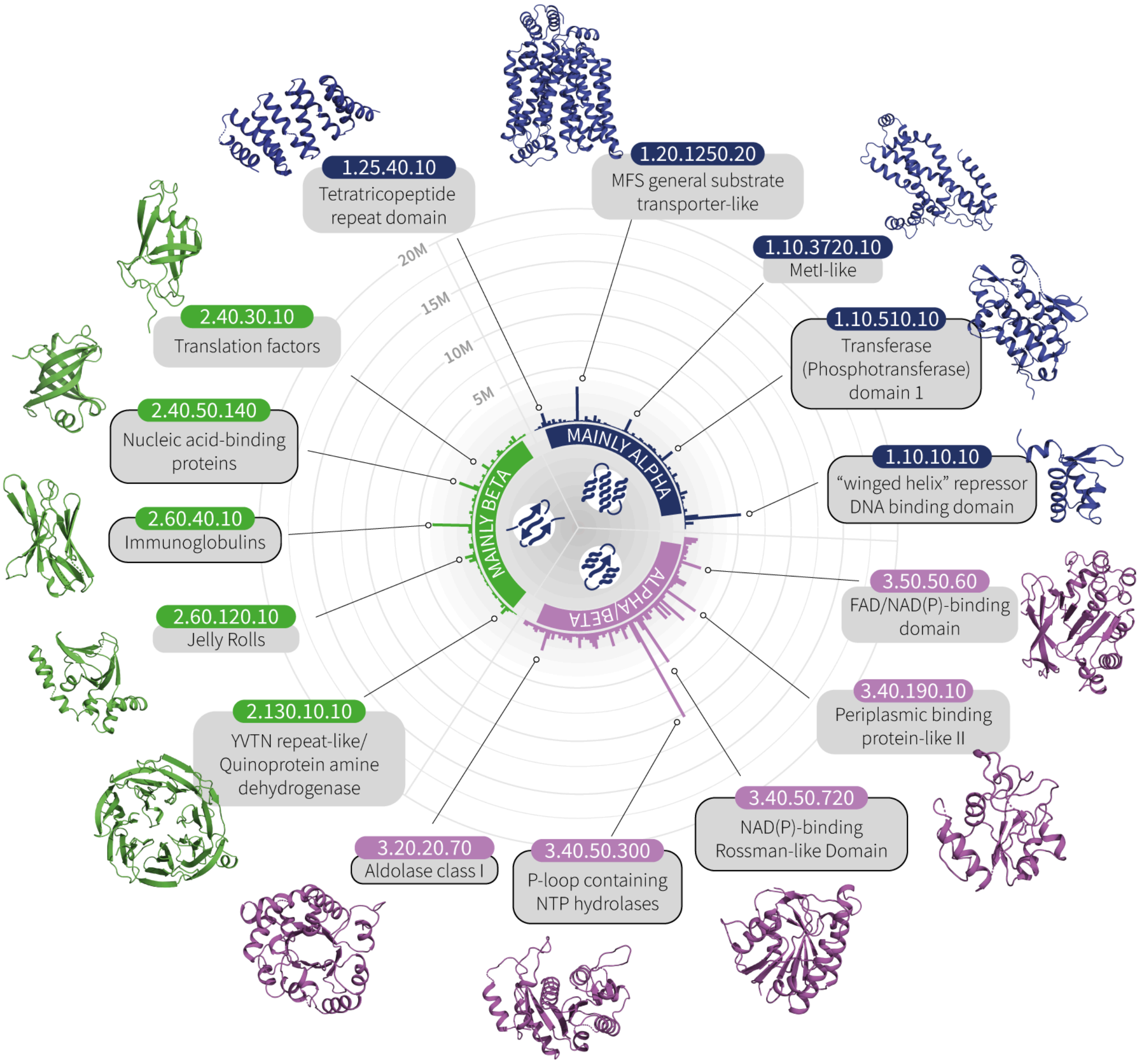

**Supplementary Figure 9. Most abundant superfamilies identified in TED-100.** The top 5 superfamilies per CATH class are shown, along with representative domain structures from CATH. Superfamilies that are part of the top 5 most abundant in CATH are outlined in black (Supp. Figure 10).

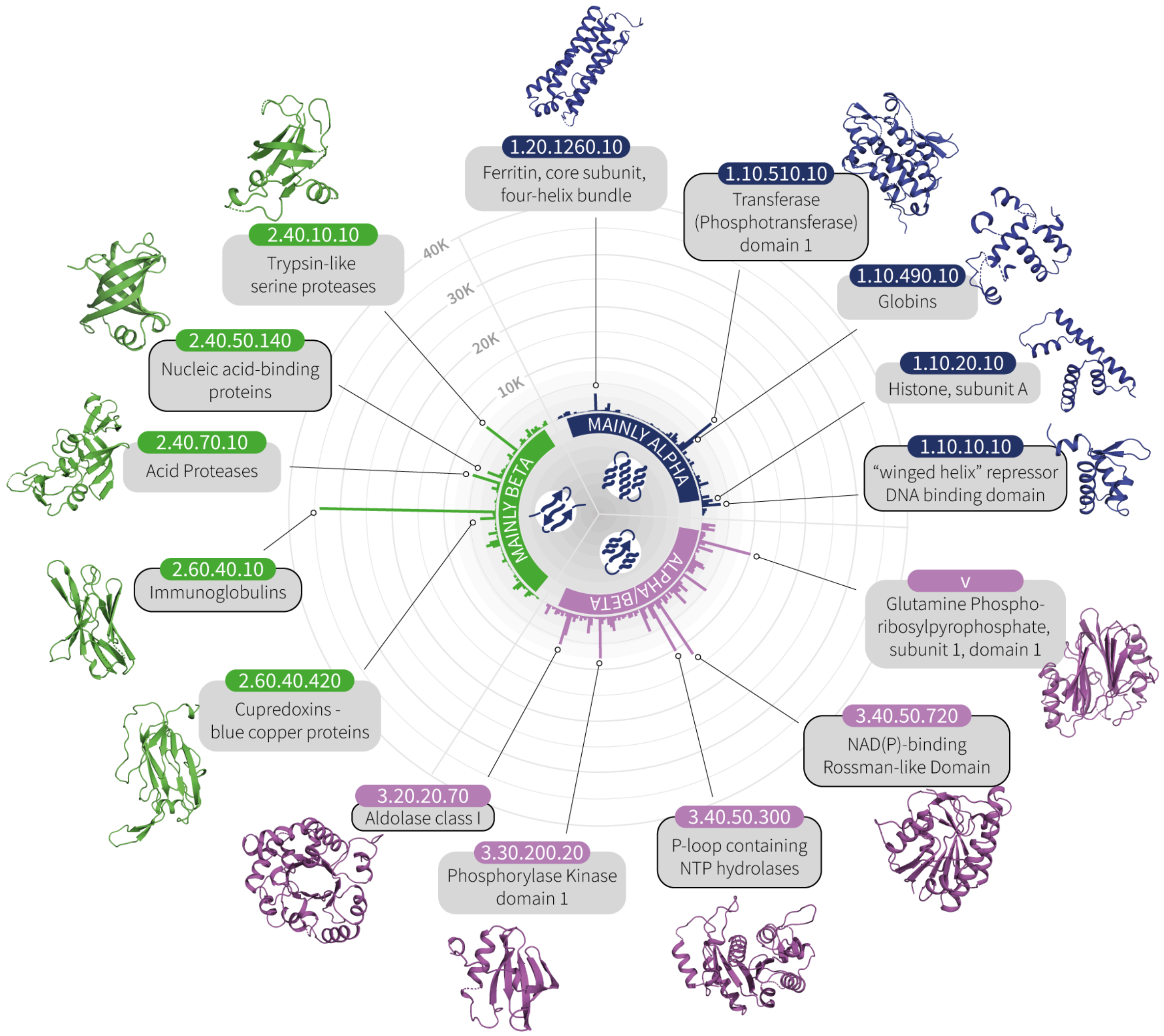

**Supplementary Figure 10. Most abundant superfamilies in CATH.** The top 5 superfamilies per CATH class are shown, along with representative domain structures from CATH. Superfamilies that are part of the top 5 most abundant in TED-100 are outlined in black (Supp. Figure 9).

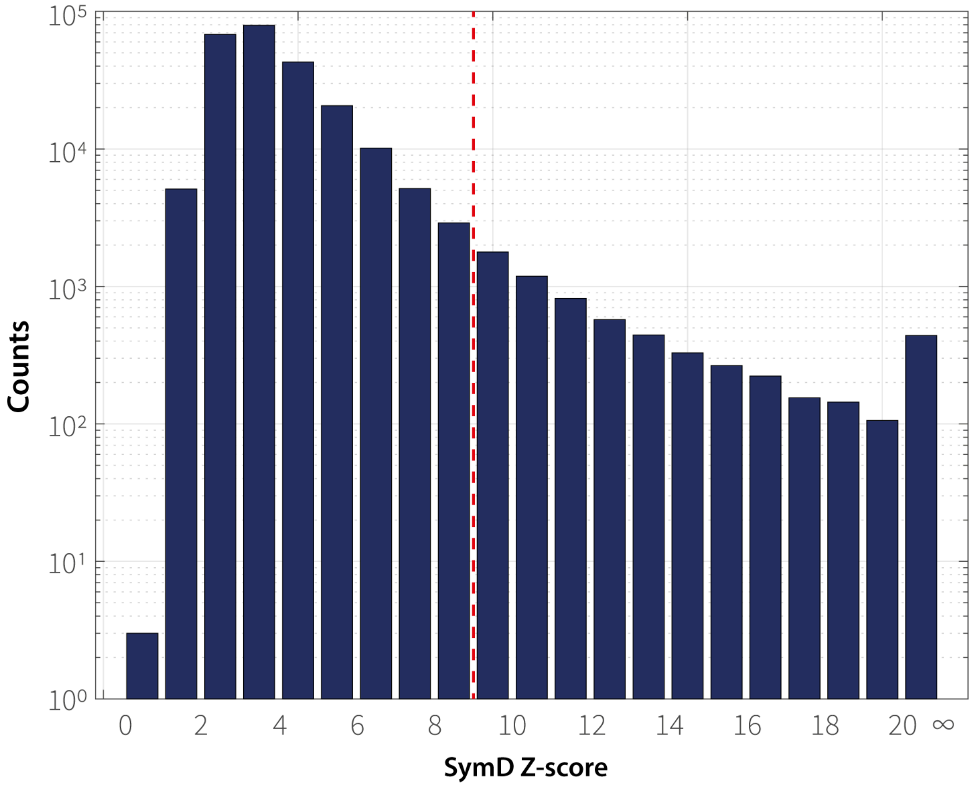

**Supplementary Figure 11. Distribution of symmetry Z-scores calculated on candidate novel domains.** Z-scores are calculated using the SymD program on TED domain structures directly (n=240,674). The red line demarcates the cutoff of 9 used for filtering high-symmetry domains.

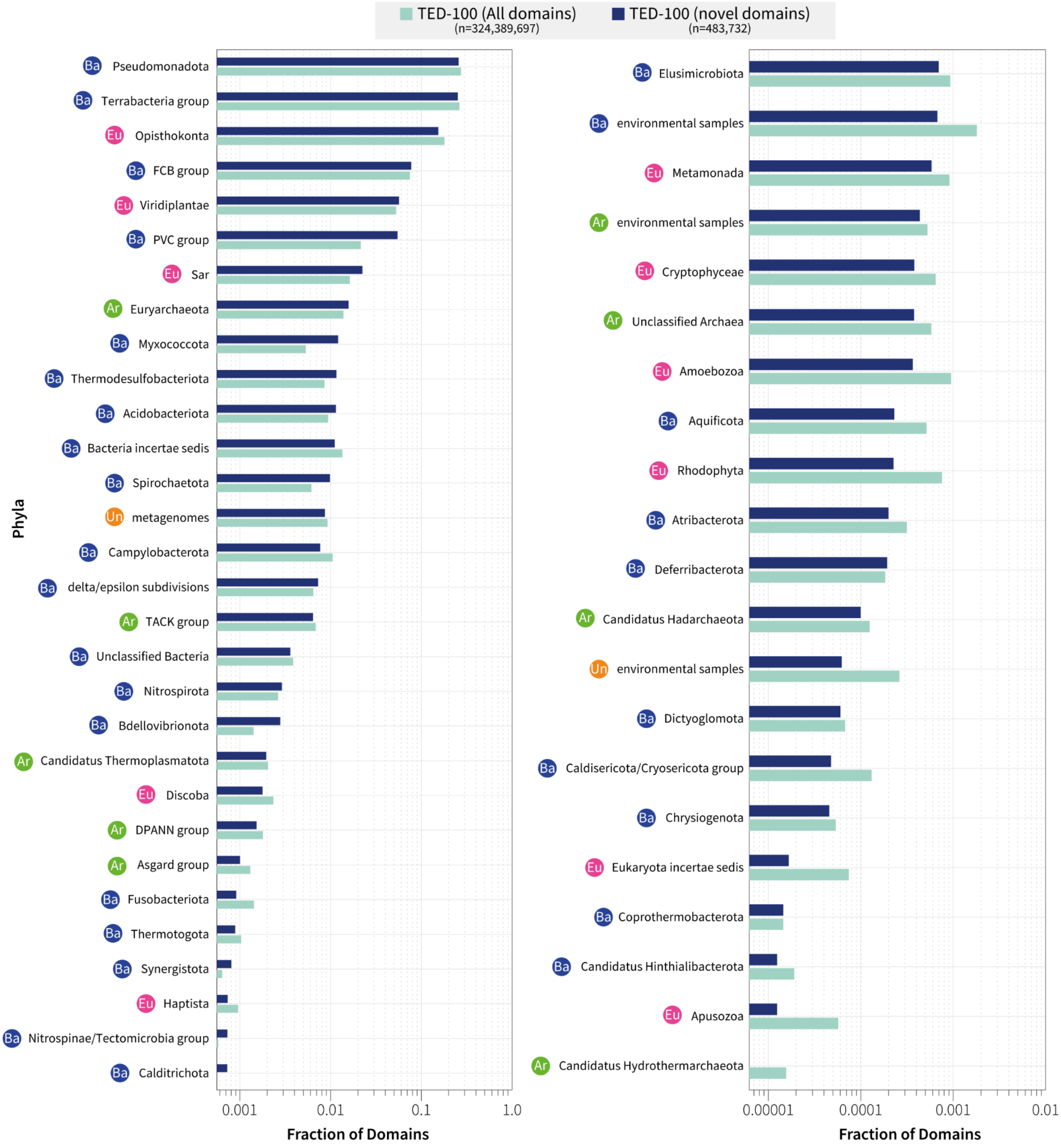

**Supplementary Figure 12**. **Comparison of taxonomic distribution of identified novel domains against baseline proportions in TED-100.** Bar charts compare the fraction of domains found in each phyla for novel domains (navy) and all domains in TED-100 (light blue). Phyla are ranked according to the novel domain fraction found. Labels next to each phylum denote the superkingdom: bacteria (Ba), eukaryota (Eu), archaea (Ar) and unclassified (Un).

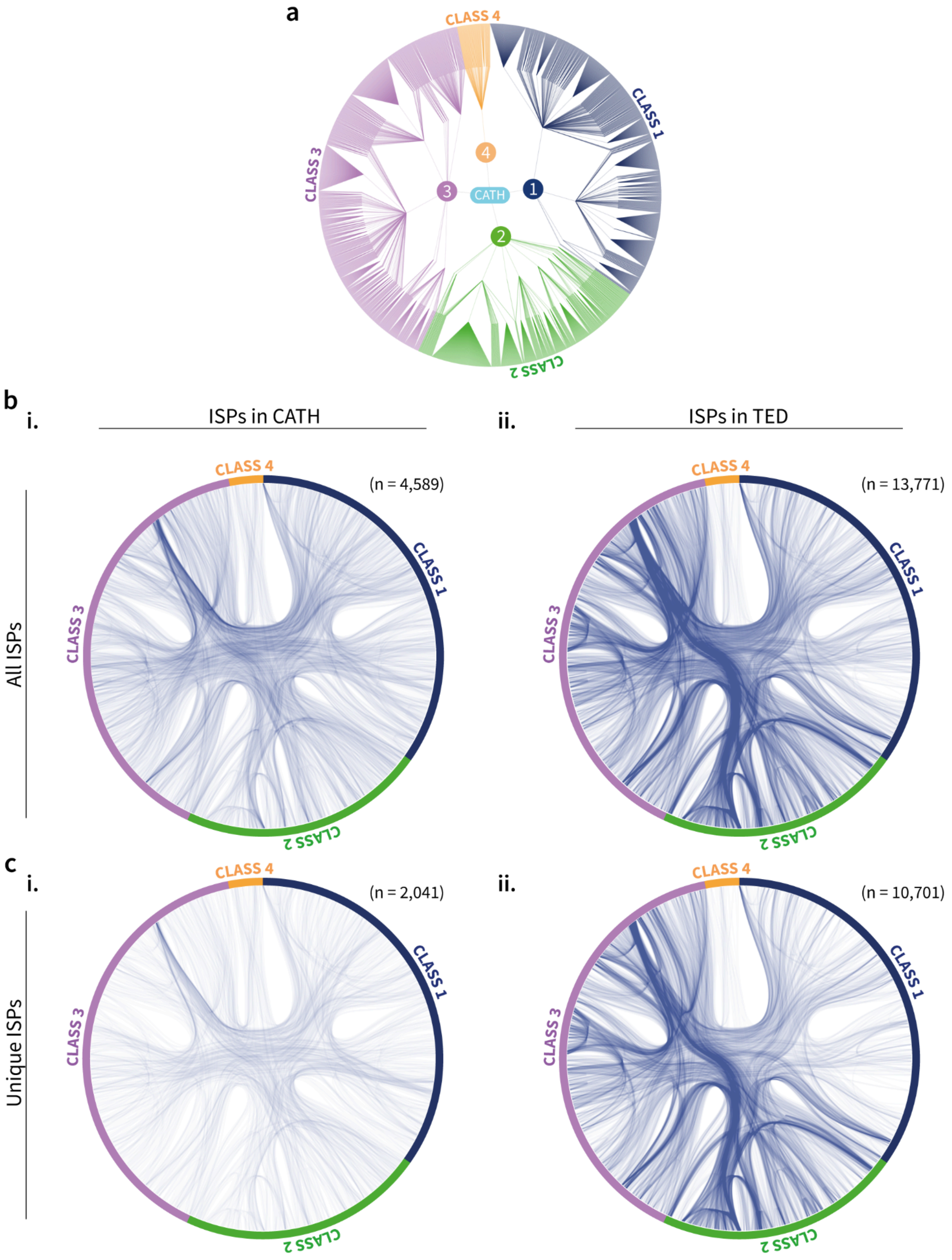

**Supplementary Figure 13.** (**a**)**.** The CATH hierarchy (classes 1-4 only) is represented as a circular dendrogram with a virtual root node for the 4 classes (labelled 'CATH'). Edges are coloured according to CATH class. Paths drawn in the hierarchical edge bundling diagrams in (b) and (c) follow the layout of this dendrogram smoothly when connecting superfamilies, which are the leaf nodes on the outer edge. Hierarchical edge bundling diagrams representing (**b**) all ISPs and (**c**) ISPs unique to either set, for i. CATH, and ii. TED.

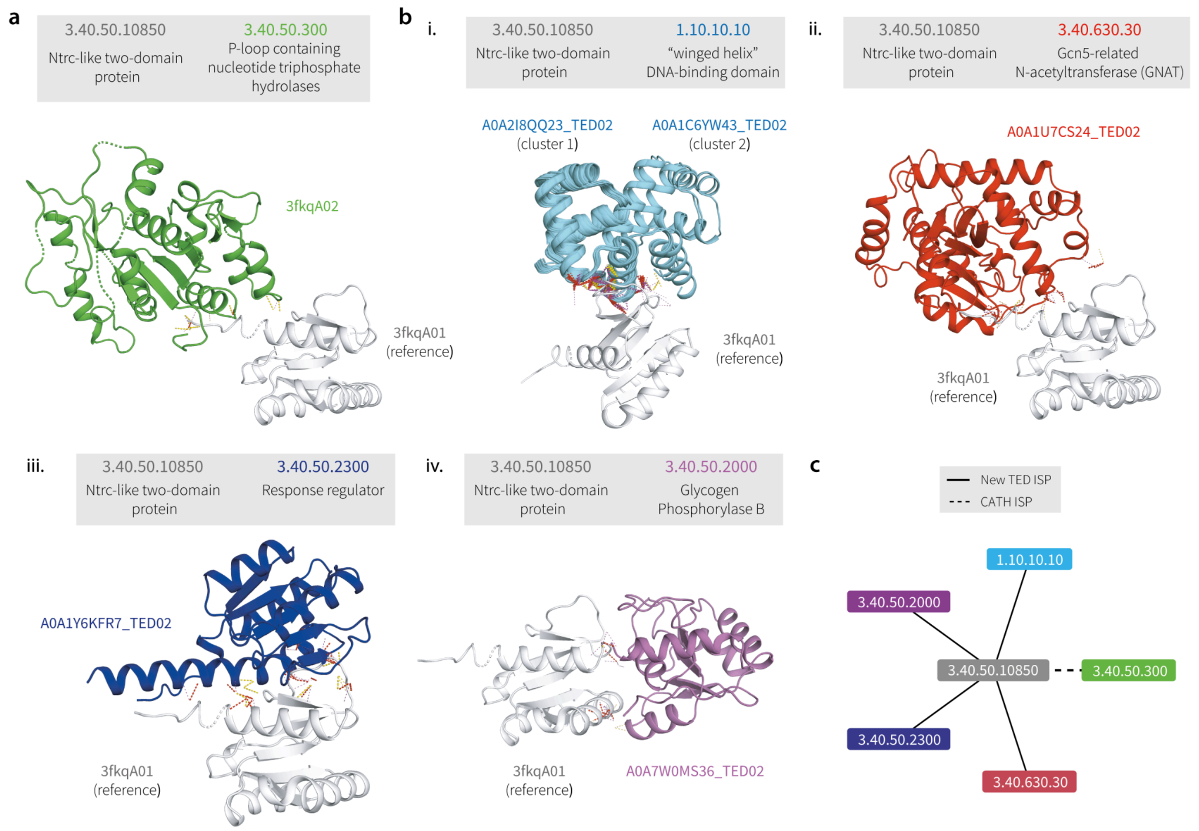

**Supplementary Figure 14. Interactions for CATH superfamily 3.40.50.10850.** (**a**) In the PDB, the NtrC-like protein domain (superfamily 3.40.50.10850) is only found interacting with a single P-loop-containing nucleotide triphosphate hydrolase family (superfamily 3.40.50.300). (**b**) In TED, superfamily 3.40.50.10850 can be seen interacting with four additional CATH superfamilies - i. 1.10.10.10 ("winged-helix" DNA-binding domain), ii. 3.40.630.30 (Gcn5-related N-acetylransferase), iii. 3.40.50.2300 (response regulator) and iii. 3.40.50.2000 (glycogen phosphorylase B) superfamilies. (**c**) Network graph showing existing interactions seen in CATH (PDB) and new interactions observed in TED for superfamily 3.40.50.10850.

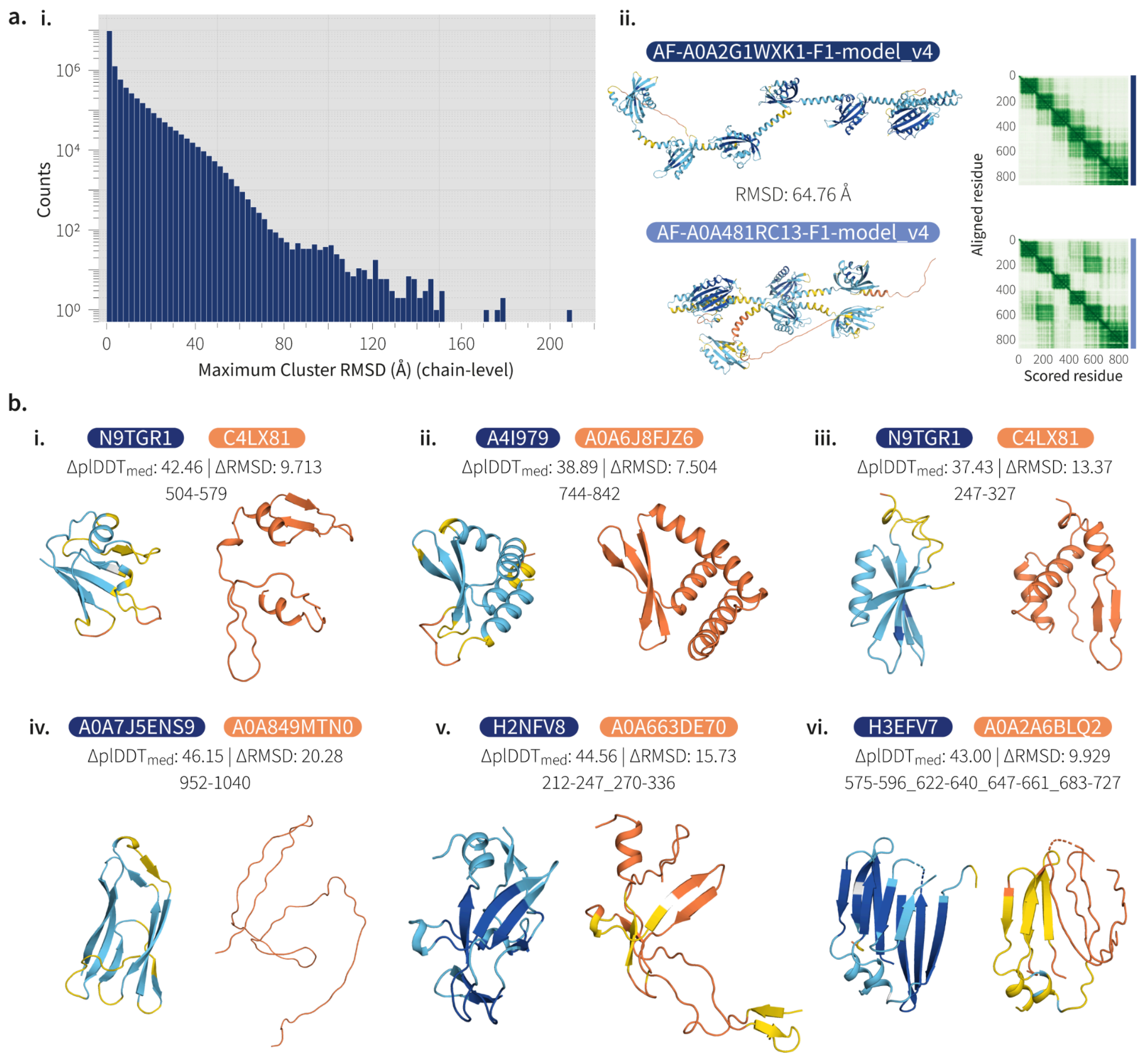

**Supplementary Figure 15. Structural diversity across identical sequences in the AFDB.** (**a**) i. Distribution of maximum RMSD observed across structures of identical sequences (n=13,175,417). For each identical sequence cluster, the maximum RMSD observed across pairs of structures is recorded. Dataset comprises all TED-redundant targets (38,944,835 chains across 13,175,417 unique sequences). ii. Example of a pair of structures with identical sequences (full chain). Dark areas in PAE maps represent high confidence in the positioning between pairs of residues. Differences can be seen in domain packing as well as in the PAE map produced by AF2. AF2 has greater confidence in the packing of domains 2 and 5 in the bottom structure compared to the top. (**b**) i-vi. Examples of structural diversity in domains of identical sequence models. In each example, the difference in median domain pIDDT as well as RMSD following TMalign is shown. Domain structures represent the exact same residue ranges in two models with identical full chain sequences. Colouration follows pIDDT confidence bins as per the AFDB (dark blue/very high: pIDDT >= 90, blue/high: 90 > pIDDT >= 70, yellow/low: 70 > pIDDT >= 50 and orange/very low: pIDDT < 50).

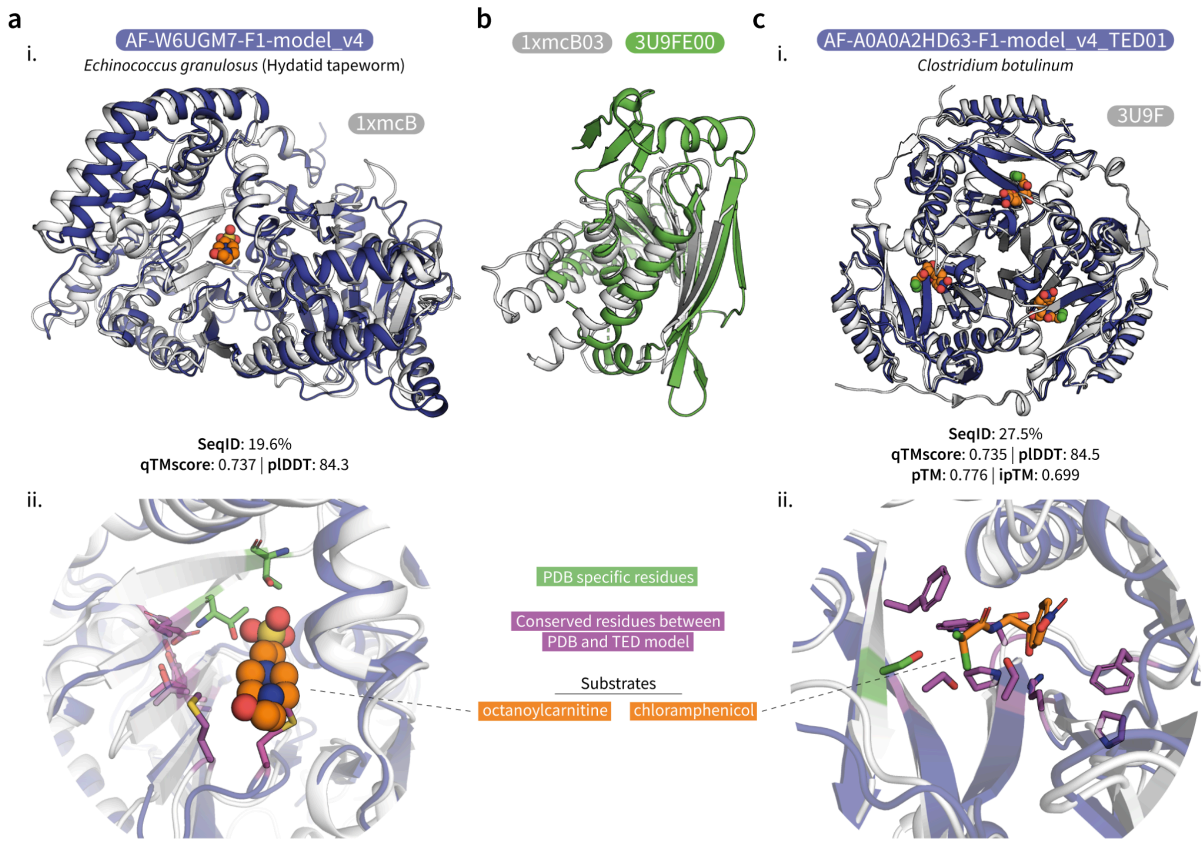

**Supplementary Figure 16. Structural insights into differences and commonalities among CoA-dependent acyltransferases.** (**a**) i. Superposition of AF-W6UGM7-F1-model_v4 and 1xmcB. ii. Substrate binding pocket of (a) with substrate (orange), conserved residues unique to PDB:1xmc (green) and shared between PDB:1xmc and TED monomer (magenta). (**b**) Superposition of CATH structural domain representatives clustered at 5Å for homotrimers (CATH: 3u9FE00) and monomers (1xmCB03). (**c**) i. Superposition of 3u9F and a modelled trimer of AF-A0A0A2HD63-F1-model_v4_TED01. ii. Substrate binding pocket of (c) with substrate (orange), conserved residues unique to PDB:3u9F (green) and shared between 3u9F and TED trimer (magenta).

Present in most taxonomic branches of the Tree of Life, the chloramphenicol acetyltransferase-like domain superfamily in CATH (3.30.559.10) adopts an αβα sandwich fold. The superfamily comprises a wide range of enzymes that function as CoA-dependent acyltransferases, with chloramphenicol acetyltransferases among others. Members of the superfamily share a conserved HXXXD motif in their active site and their oligomerization state can vary dramatically, from the more preponderant 2-domain single monomer **(a)**, to a more complex homo-trimer (**c**). Their oligomeric modes are fundamental to their catalytic functions. For example, carnitine octanoyltransferase functions as a monomer and comprises a N-terminal and a C-terminal domain which together form a substrate binding pocket [PDB ID: 1xmc, **a**], whereas chloramphenicol acetyltransferases adopt a homo-trimeric assembly for substrate-binding and catalytic activity[10] [PDB: 3u9F, **c**]. Ubiquitous and with a known biochemistry, chloramphenicol acetyltransferases play a critical role in antibiotic resistance in pathogens and are prime targets for drug repurposing aimed at untreated pathogens of interest in health and agriculture.

TED vastly expands the structural coverage of the superfamily, from 253 domains in CATH to 228,867 in TED (Supp. Methods), aiding knowledge transfer of conserved residues and oligomerization states between remote homologs. Low sequence identity can be bridged by structure, enabling the identification of very remote homologs across TED using matching by structure instead of sequence. In panels a) and d), a remote homolog (AF-W6UGM7-F1-model_v4_TED03; Carnitine O-palmitoyltransferase; EC 2.3.1.21) of mouse carnitine octanoyltransferase (PDB:1xmcB003; carnitine octanoyltransferase; EC: 2.3.1.137) can be detected in hydatid tapeworm (qTMscore=0.737), a domestic pathogen by structure matching despite having a low sequence similarity (19.6%). By clustering highly structurally similar TED relatives (Supp. Methods), we could predict highly conserved residues for the tapeworm domain. The majority of these occur in a pocket which superimposes with the known substrate binding region in the experimental structure and furthermore the majority of the predicted conserved residues in the tapeworm domain superimpose well with those in the mouse domain. Despite this, structural analysis shows that the PDB structure has an insertion of two additional beta strands in the substrate binding pocket, which possess conserved residues unique to the PDB domain (shown in blue in panel a-ii)) suggesting a role in the distinct substrate specificity (2.3.1.137 rather than EC 2.3.1.21) of this enzyme.

Panel c) show superposition of a structural homolog in AFDB (AF-A0A0A2HD63-F1-model_v4_TED01), of a known chloramphenicol acetyltransferase (3u9F in the PDB). The AFDB homolog is from *Clostridium botulinum*, a pathogenic bacteria causing botulism in humans and animals. Whilst the sequence similarity is below 30%, the TED domain matches the PDB with a TM-Score 0.82. Following modelling of the trimeric oligomer of this protein we observe a good superposition with the trimer of PDB structure. We also see that highly conserved residues in the TED domain (identified from related structures in the AFDB) superimpose well with conserved residues in the PDB domain in the Chloramphenicol-binding site (see figure e), supporting similar substrate binding.

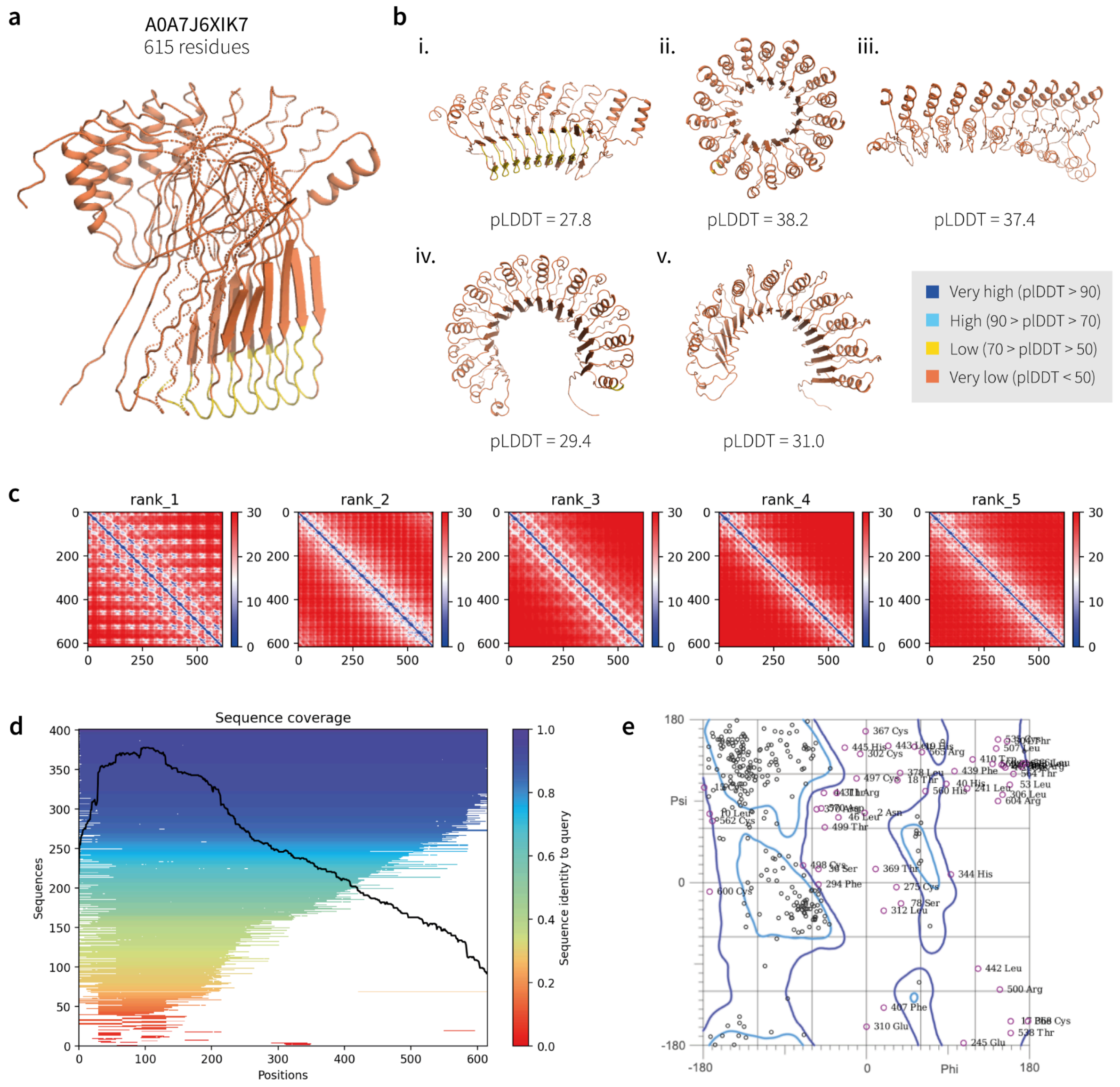

**Supplementary Figure 17. Example of a problematic structure in the AFDB.** (**a**) The AFDB model of Uniprot entry A0A7J6XIK7 from *Trypanosoma cruzi* is a low plDDT structure with a high proportion of clashes. (**b**) Re-prediction of the structure using Colabfold generates several visually plausible structures with high symmetry and repetition. All models are predicted with very low plDDT. (**c**) PAE output for each of the five structures is shown in panel (b). Blue colouration indicates high confidence between the position of residue pairs. (**d**) Sequence coverage of the MSA generated during remodelling in (b). (**e**) Molprobity Ramachandran plot of model shown in (a). Text-labelled residues have torsion angles outside of allowed regions.

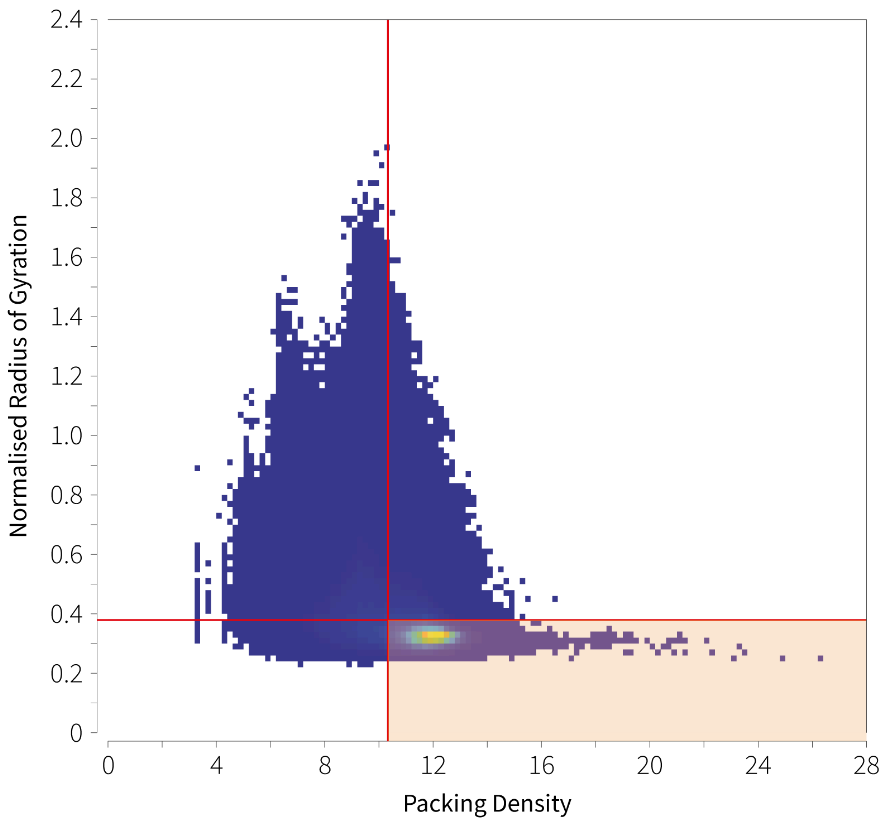

**Supplementary Figure 18. Globularity assessment of domains in TED-100.** Globularity of 324m TED-100 domains were assessed with CATH-AlphaFlow using packing density and normalised Rg metrics as described in Bordin et al.[1]. To determine suitable cutoffs for identifying globular domains, thresholds were calculated as the 5th percentile of each metric on the subset of TED-100 domains which were assigned high-confidence CATH superfamily labels (193m domains). Packing density and normalised Rg cutoffs were 10.333 (vertical red line) and 0.356 (horizontal red line) respectively. Colour gradient on bivariate histogram represents the density of data, where yellow is highest, and navy is lowest. Shaded red box encapsulates 244,166,721 domains deemed globular.

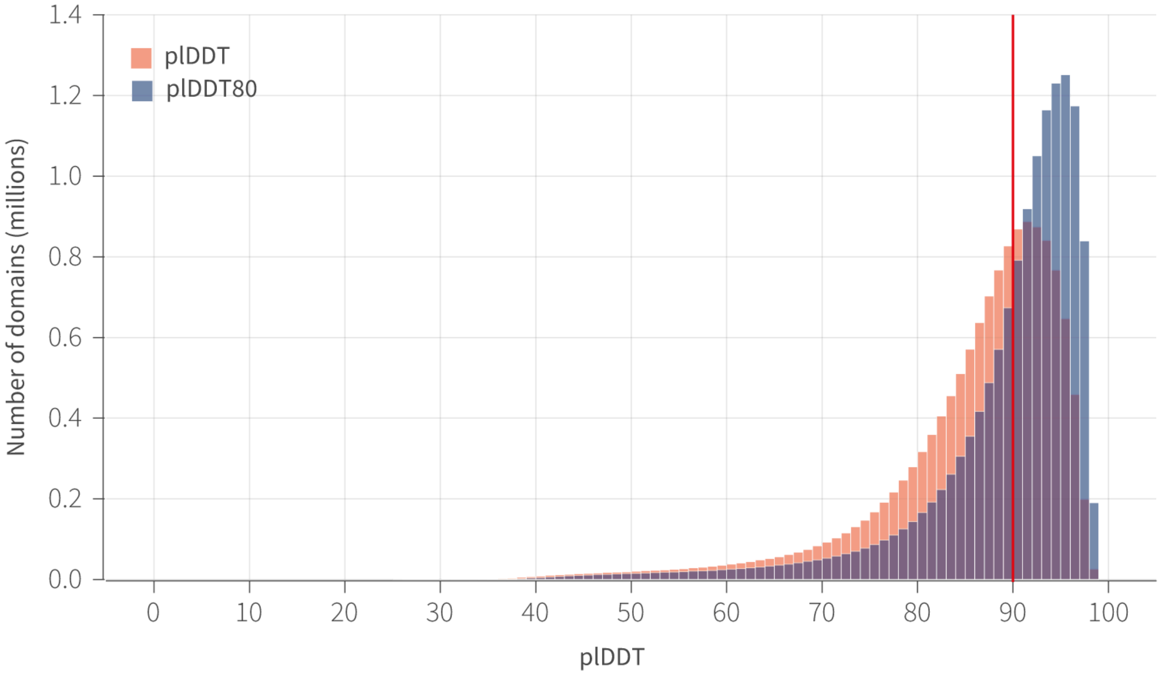

**Supplementary Figure 19. Comparison of plDDT metrics for model quality filtering.** Histograms shown represent the plDDT and $\text{plDDT}_{80}$ metrics used for model quality filtering. Data comprises 13,820,550 domains which pass globularity and nSSE filters. plDDT is calculated as the average plDDT of all residues in the domain while $\text{plDDT}_{80}$ takes into account the average residue plDDT over the highest 80% of residues. A $\text{plDDT}_{80}$ >= 90 is used for filtering domains before the novel domain identification workflow, resulting in 8,612,318 domains.

### References

1. Bordin, N. *et al.* AlphaFold2 reveals commonalities and novelties in protein structure space for 21 model organisms. *Commun Biol* **6**, 160 (2023).
2. Zhang, Y. & Skolnick, J. TM-align: a protein structure alignment algorithm based on the TM-score. *Nucleic Acids Res.* **33**, 2302–2309 (2005).
3. Frishman, D. & Argos, P. Knowledge-based protein secondary structure assignment. *Proteins* **23**, 566–579 (1995).
4. Zhou, N. *et al.* The CAFA challenge reports improved protein function prediction and new functional annotations for hundreds of genes through experimental screens. *Genome Biol.* **20**, 244 (2019).
5. UniProt Consortium. UniProt: the Universal Protein Knowledgebase in 2023. *Nucleic Acids Res.* **51**, D523–D531 (2023).
6. Orengo, C. A. & Taylor, W. R. SSAP: sequential structure alignment program for protein structure comparison. *Methods Enzymol.* **266**, 617–635 (1996).
7. Laskowski, R. A., Jabłońska, J., Pravda, L., Vařeková, R. S. & Thornton, J. M. PDBsum: Structural summaries of PDB entries. *Protein Sci.* **27**, 129–134 (2018).
8. Valdar, W. S. J. Scoring residue conservation. *Proteins* **48**, 227–241 (2002).
9. Mirdita, M. *et al.* ColabFold: making protein folding accessible to all. *Nat. Methods* **19**, 679–682 (2022).
10. Biswas, T., Houghton, J. L., Garneau-Tsodikova, S. & Tsodikov, O. V. The structural basis for substrate versatility of chloramphenicol acetyltransferase CATI. *Protein Sci.* **21**, 520–530 (2012).
